# soibean: High-resolution Taxonomic Identification of Ancient Environmental DNA Using Mitochondrial Pangenome Graphs

**DOI:** 10.1101/2024.04.12.589157

**Authors:** Nicola Alexandra Vogel, Joshua Daniel Rubin, Anders Gorm Pedersen, Peter Wad Sackett, Mikkel Winther Pedersen, Gabriel Renaud

## Abstract

Ancient environmental DNA (aeDNA) is becoming a powerful tool to gain insights about past ecosystems. However, several methodological challenges remain, particularly for classifying the DNA to species level and conducting phylogenetic placement. Current methods, primarily tailored for modern datasets, fail to capture several idiosyncrasies of aeDNA, including species mixtures from closely related species and ancestral divergence. We introduce `soibean`, a novel tool that utilises pangenomic graphs for identifying species from ancient environmental mitochondrial reads. It outperforms existing methods in accurately identifying species from multiple sources within a sample, enhancing phylogenetic analysis for aeDNA. `soibean` employs a damage-aware likelihood model for precise identification at low-coverage with high damage rate, demonstrating effectiveness through simulated data tests and empirical validation. Notably, our method uncovered new empirical results in published datasets, including using porpoise whales as food in a Mesolithic community in Sweden, demonstrating its potential to reveal previously unrecognised findings in aeDNA studies.

## 1 Introduction

Ancient DNA (aDNA) provides a crucial window into identifying eukaryotic species from ancient remains by giving additional insight into archaeological and paleontological findings. However, fossils and other macroscopic remains are merely partial sources of information. Recently, ancient environmental DNA (aeDNA) has changed our understanding of past environments and species compositions in both time and space. Throughout an organism’s lifetime, it leaves genetic traces in the environment, in deposits such as sediment or permafrost [15, 70, 20, 32, 69]. The extraction and amplification of aeDNA allow for a widely distributed exploration of past ecological environments and populations from chosen sample sites [23, 13, 8, 7, 40, 71, 41]. However, analysing aDNA from the environment and bones poses many challenges. Firstly, aDNA is characterised by being highly fragmented [39, 19] and modified by chemical damage such as deamination patterns (resulting in C to T and G to A substitutions) [4, 46]. This causes changes in the similarity to the reference genomes used for taxonomic identification [45, 34]. In addition, aDNA analysis must also consider the evolutionary processes, hence genetic distance between the organism and the reference genome and even the absence of a reference genome [44, 54].

aeDNA inherits all the challenges associated with aDNA, while also presenting the added complexity of being a mixture of DNA from various sources. The accurate taxonomic classification of aeDNA fragments is greatly influenced by the relative abundance of DNA from each contributing source. Therefore, taxonomic classification can be either of low specificity (e.g. class, order, family) or high specificity (e.g. species, subspecies). With lower abundance, achieving a high taxonomic specificity is often more challenging due to a lack of unique genetic identifiers [58]. In mitochondrial aeDNA analysis, results are often summarised at a lower taxonomic specificity [59, 26] when using standard classification methods like a naive lowest common ancestor (LCA) algorithm [2, 22, 68]. A newer classification tool, euka, also classifies at lower taxonomic resolutions [65]. This classification method aids a confident validation of identified taxa via damage pattern estimation [38] or estimation of breadth and depth of coverage due to an increased amount of aeDNA fragments for a given taxon.

One method for high-resolution taxonomic assignment was proposed with `HAYSTAC` [6]. `HAYSTAC` provides verification filters (e.g. likelihood filter, coverage evenness filter) for accurate species detection. Its all-versus-all mapping approach with Bowtie2 [27] considers all possible mapping positions, including those within highly conserved regions across species, which are usually ignored due to their inability to discriminate at the species level. For aeDNA analysis, these regions can be useful due to the sparsity of the data. We use `HAYSTAC` as our baseline model as it allows us to provide a user-built database. However, it does not account for private mutations or place samples within a phylogenetic reference. This limitation makes it challenging to identify the ancestral species.

Another method to more confidently assign classifications to a species or lower is phylogenetic placement, in which a consensus is called from the extracted fragments and placed on a phylogenetic tree based on sequence similarity [3, 11, 63]. However, aeDNA data is often too low coverage to reliably call a consensus and, therefore, unfitted for phylogenetic placement. Furthermore, this problem becomes intractable if multiple species from the same genus (e.g., Arctic, Mountain, and Snowshoe hares) [67] or closely related species [42] exist.

To our knowledge, the only tool for species detection in low-coverage aDNA data is `pathPhynder` [33]. `pathPhynder` considers unique SNPs to identify the most likely species. `pathPhynder` considers all derived and ancestral SNPs on a phylogenetic tree and is, therefore, able to infer a potential ancestral state of a species, making it extremely valuable for aDNA analysis [26]. However, `pathPhynder` is limited to single-source estimations. Multiple sources must be mapped beforehand and analysed individually [42], which can adversely affect abundance estimates. Moreover, `pathPhynder` does not consider insertions or deletions in alignments, potentially discarding useful information.

We introduce `soibean`, a new subcommand of `vgan` (https://github.com/grenaud/vgan) for high-resolution taxonomic placement of aeDNA using mitochondrial pangenome graphs in conjunction with Bayesian inference methods. Pangenome graphs are reference data structures that mitigate reference bias by representing multiple genomic sequences simultaneously [10, 34, 57]. `soibean`’s input is a FASTQ file consisting of aDNA fragments that have been previously classified to a lower taxonomic specificity, such as family level (e.g. with an LCA tool or euka). `soibean` then deconvolves reads into each contributing source at the species level and subsequently places them in their phylogenetic context. Our algorithm works as follows: i) We align the aeDNA fragments to a curated and quality-controlled database of 326 arthropodic and tetrapodic taxa, including reconstructed ancestral states, allowing for variation unseen in modern reference genomes. ii) `soibean` then uses Markov Chain Monte Carlo (MCMC) sampling to estimate the most likely placement on a phylogenetic tree branch and the relative abundances of each source, allowing robust identification from *∼*50 fragments. iii) `soibean`’s results provide credible intervals and diagnostic metrics for all parameters of each source. Crucially, we can identify ancestral states and visualise confidence in phylogenetic placements. If a source has scarce data, either due to low relative abundance or simply low coverage for the taxon, our algorithm displays the uncertainty as the MCMC chains will sample widely across the tree branch. Runtimes depend on the number of iterations and input reads but ranged from 0.5 to 450 hours on example data we analysed.

This manuscript demonstrates `soibean`’s specificity and sensitivity on simulated datasets (one to four different sources on five different taxa) before highlighting its consistency with empirical data. Lastly, we showcase `soibean`’s ability to discover novel results from previously published datasets, including discovering harbour porpoise as a food source for a Mesolithic community in Sweden.

## 2 Results

To generate our results, we used `soibean`’s default settings (commands for the simulated datasets in Supplementary Subsection 1.3 and empirical datasets in Supplementary Subsection 1.6) on different pangenome graphs from `soibean`’s database (construction pipeline illustrated in Supplementary Figure 1). `soibean`’s general workflow starts with an initial estimation of the number of sources present in the sample, followed by the MCMC estimation of each source most likely placement on the phylogenetic tree and their relative abundance, and ends with MCMC diagnostics and output visualisation (see Figure 1). Detailed explanations and methodological descriptions of the various aspects of our algorithm are found in the Method Section 4. `soibean` runtime depends on the size of the dataset and the number of MCMC iterations. An average time consumption chart for users is presented in Supplementary Figure 27, with a comprehensive computational analysis available in Supplementary Subsection 1.1.

**Figure 1:**
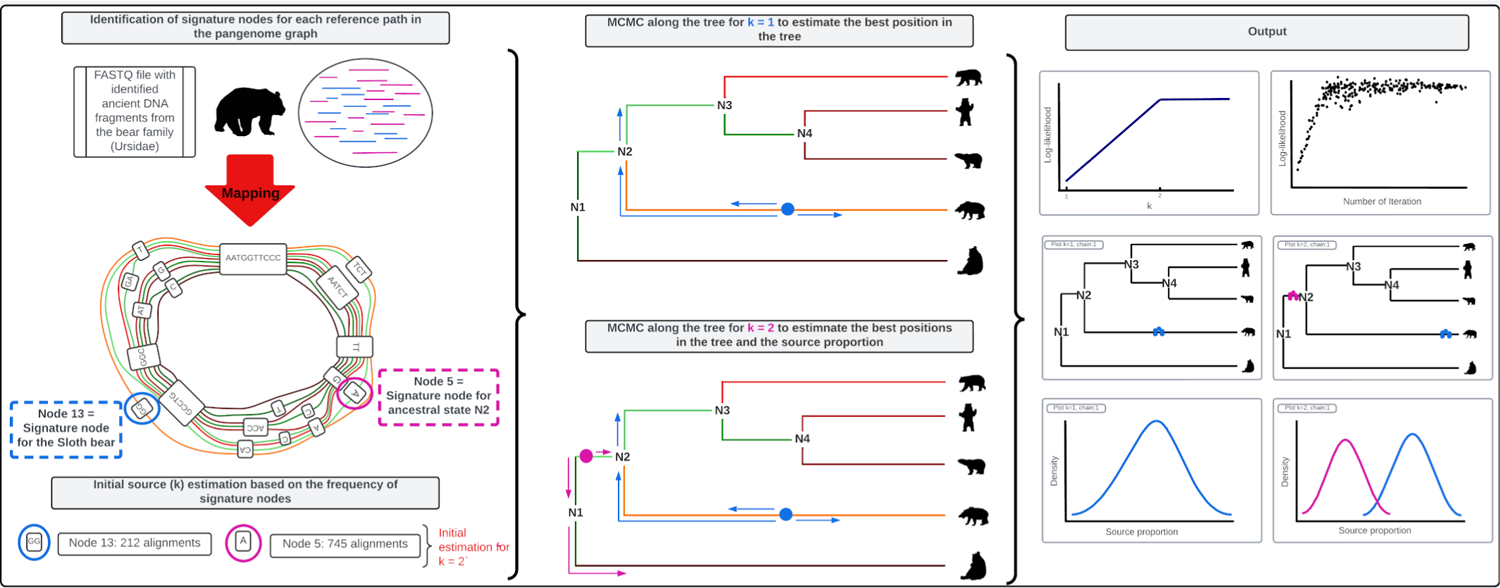
soibean’s main workflow starts by mapping a filtered FASTQ file against the selected taxon graph. The alignment is analyzed, and an initial source estimation based on signature note-set frequencies is calculated. `soibean` runs an MCMC algorithm from 1 to the number of estimated sources and calculates a source proportion and a branch position for each source based on a maximum likelihood function. The MCMC diagnostics provide statistics about the most likely number of sources, their proportion, and branch position. `soibean` provides extensive plotting scripts to visualise its results.

### 2.1 Simulations

To test `soibean`, we simulated six different datasets. For the first dataset, we simulated a single source from an ancestral state sequence from the family of bears (Ursidae). From the same family, we simulated another three datasets for two-source samples. The first two-source mixture contains two closely related bear species (*∼* 98.8% genome similarity), the second mixture contains two less closely related bear species (*∼* 93.1%) and lastly, a mix of two divergent bear species (*∼* 83.4%). We simulated a three-source dataset using a family of winged insects (Saturniidae), sampling from two emperor moth species and an ancestral state. The last dataset was to simulate a four-source dataset, where we used the family of earless seals (Phocidae). We simulated reads from four species of the same genus (*Phoca*). Simulations were created with `gargammel` [49], where each dataset has a fragment length distribution following a log-normal distribution with *µ* = 3.7344 and a *σ* = 0.35 as commonly seen in aDNA studies and deamination rates taken from Günther et al. [14]. We merged the simulated reads with leeHom using ancient parameters [48]. All simulated datasets are in our provided test data https://github.com/nicolaavogel/soibeanDatabase.

All simulations were used for benchmarking against `HAYSTAC` as our baseline model. We additionally compared our single-source simulations with pathPynder. Details can be found in Supplementary Subsection 1.2 and the commands used in Supplementary Subsection 1.3.

#### 2.1.1 Single-source

Our single-source sample is simulated from the ancestral state N4 of the bear family (Ursidae). To show `soibean`’s robustness, we downsampled the data from *∼*1.3*X* coverage to *∼*0.026*X* coverage (see Figure 2). Figure 2 shows the complete phylogenetic tree for the family of bears on the top. For each downsampled coverage, we show the MCMC’s trace plot on the left side, which has a dot for every proposed move log-likelihood and demonstrates the initial finding of the correct tree branch and the optimal exploration of the parameter space by using independent sampling. The right side shows the zoomed-in portion of the phylogenetic tree, with every accepted MCMC move as a red dot. It can be seen how locations close to the true node are sampled and how the uncertainty about the location increases for lower coverage (red and yellow dots cover a larger area surrounding the true node). Figure 2 demonstrates `soibean`’s accuracy down to *∼*0.13*X* coverage, corresponding to *∼* 50 aDNA fragments aligned to the mitochondrial genome. We can see that the certainty of branch positions decreases with lowered coverage as we accept moves across the entire branch. At *∼*0.026*X* coverage, we are unable to define the correct origin of the source. However, all accepted MCMC moves are adjacent to the true node (specifically, within the true nodes’ parent, sibling, and child branches).

**Figure 2:**
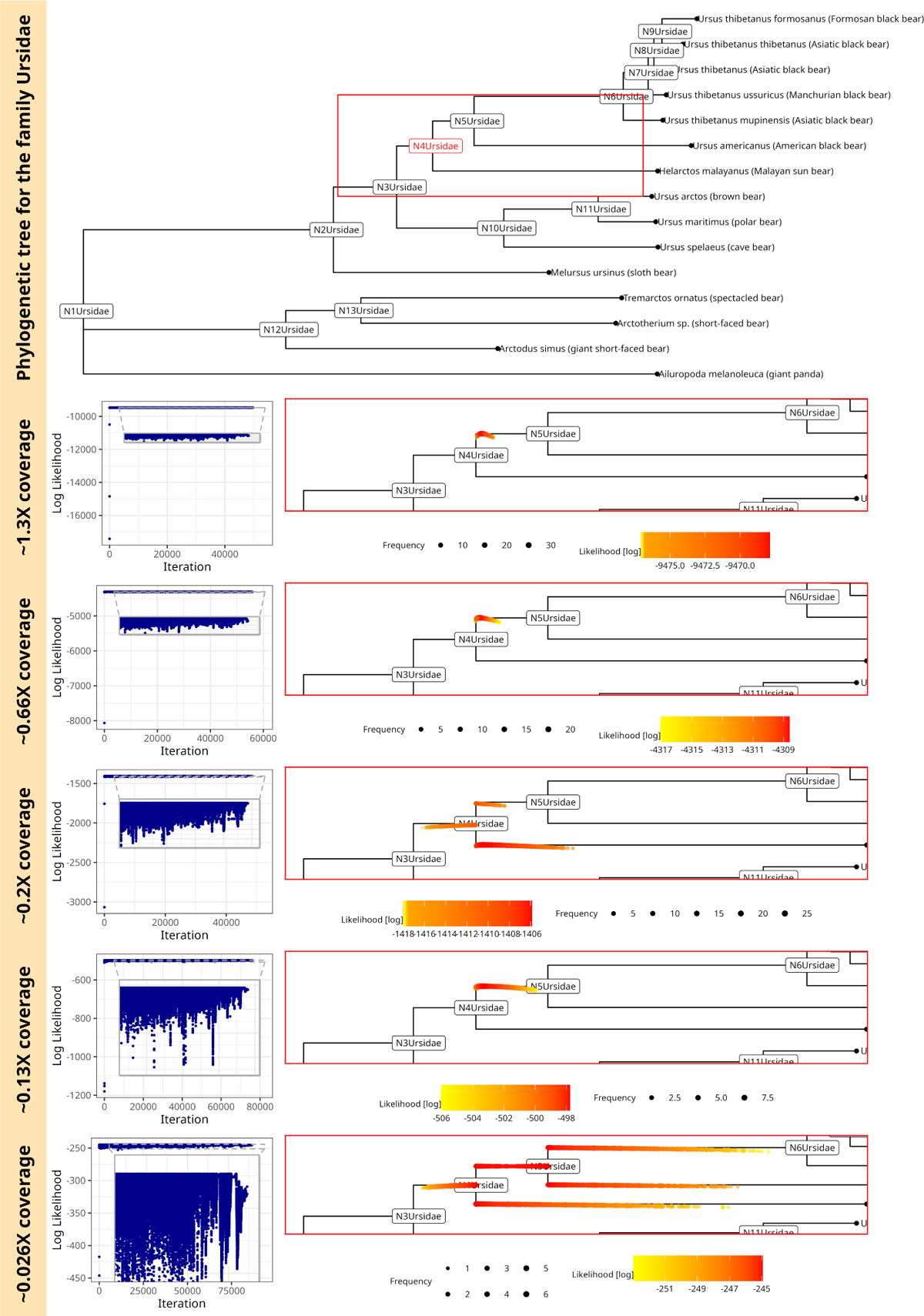
soibean results for simulated ancient fragments using a single source of DNA from the ancestral state N4 of the bear family (Ursidae). The complete phylogenetic tree for the family Ursidae marked in red is the sampled ancestral state; the red square shows the embedded zoomed-in portion of the tree in the plots below. The traceplot for the MCMC sampling and the accepted MCMC moves with their log-likelihood on the zoomed-in tree for coverage of *∼*1.3*X*(500 fragments)*, ∼*0.66*X*(250 fragments)*, ∼* 0.2*X*(75 fragments)*, ∼* 0.13*X*(50 fragments)*, ∼* 0.026*X*(10 fragments). The MCMC sampling is more uncertain with lower coverage, noted by more variable accepted moves visible in the trace plot and the zoomed-in tree plots.

Comparing `soibean`’s results to existing methods, we can observe that `pathPhynder` can accurately identify the correct ancestral source (tree node N4) down to *∼* 0.2*X* coverage with its best path method (Supplementary Figures 2, 3,4). At lower coverage, `pathPhynder` best path method predicts an incorrect source. However, source predictions stay within the targeted nodes’ parent and child nodes (Supplementary Figures 5 and 6). Additionally, we tested `pathPhynder` maximum likelihood method, which performed identical to `soibean`, correctly identifying the ancestral state N4 to *∼* 0.13*X* coverage (Supplementary Figures 7, 8, 9, 10). Only at the lowest coverage `pathPhynder` maximum likelihood method classifies incorrectly to the child (N6) of the ancestral state N4 (Supplementary Figure 11). Overall, `soibean` shows more robustness to lower coverage than `pathPhynder` best path method and performs identical to its maximum likelihood method for identifying ancestral states. `HAYSTAC` as our baseline model faces challenges identifying ancestral states despite sequences being provided (Supplementary Table 1). No ancestral state was identified at any level of coverage, likely because `HAYSTAC` was not designed to include tree topologies.

#### 2.1.2 Two Sources

For our two-source simulations, we used a mixture of two different bears: a Cave bear and a Brown bear with mitogenome similarity *∼* 93.1%. We simulated four proportions 95% *−* 5%, 85% *−* 15%, 75% *−* 25% and 55% *−* 45% with total average coverage of *∼*2.5*X* (1000 aDNA fragments). These simulations are similar to the real data seen in Pedersen et al. [42], which had a total of 740 aDNA fragments mapping for two distinct sources combined in one sample. For the mixture of the Cave and the Brown bear `soibean` identifies both sources for every simulated proportion at *∼* 2.5*X* coverage (see Figure 3). We can observe that the lower the proportion of a source (here: the Brown bear), the more uncertain the estimation of the correct position on the branch. This mirrors the observations from our single-source experiments.

**Figure 3:**
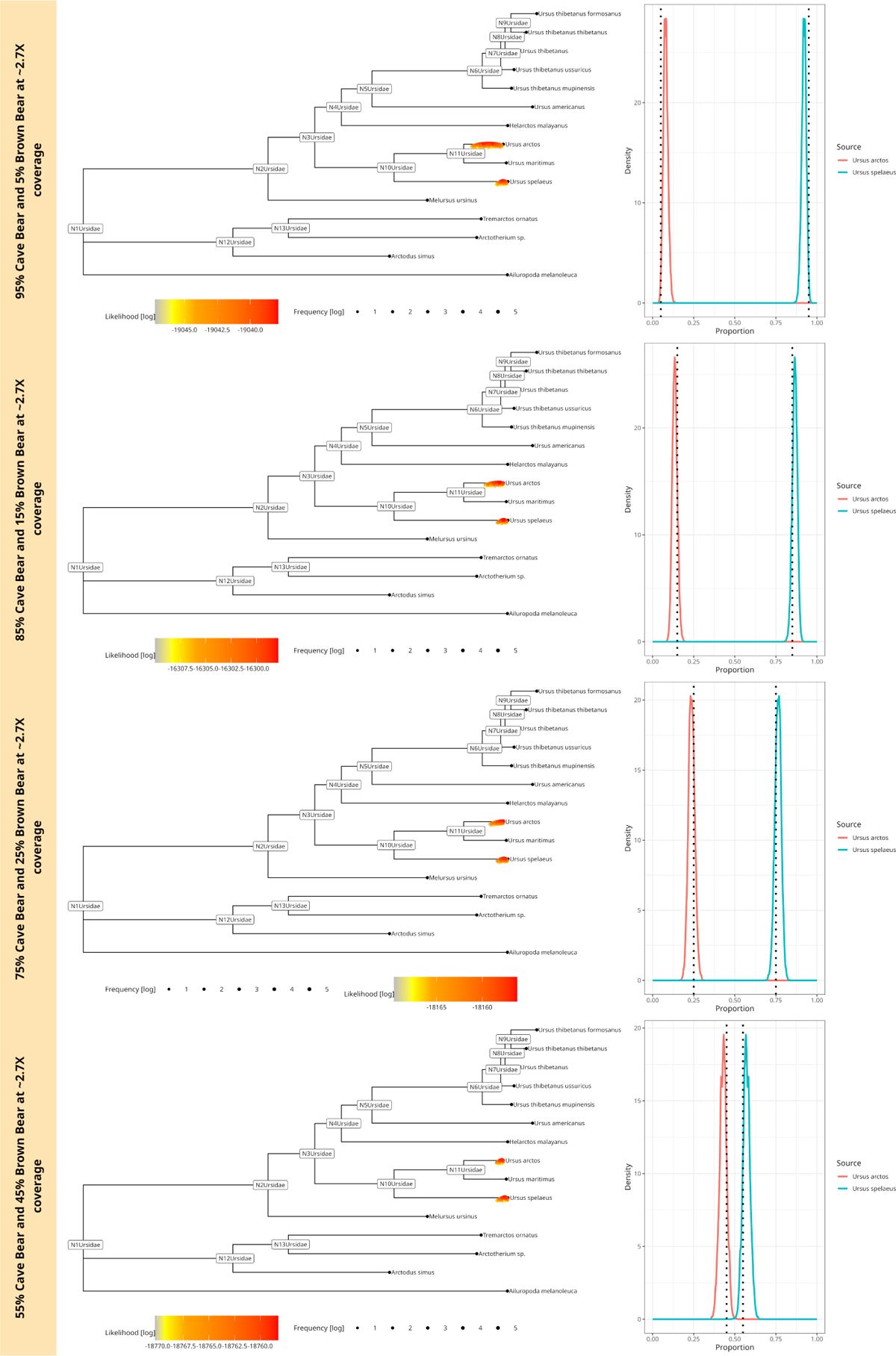
soibean results for simulated ancient fragments using a blend of two sources from two species with a similarity of 93.1% from the family bears (Ursidae) at *∼* 2*X* coverage. The plot shows four different mixtures at 55% *−* 45%, 75% *−* 25%, 85% *−* 15% and 95% *−* 5% of the Cave bear and the Brown bear. The corresponding phylogenetic trees are displayed on the left: we plotted every accepted MCMC move colored by likelihood value on the tree. The accepted moves are positioned above or below the tree, corresponding to a higher or lower likelihood value than the median, respectively. Each neighbouring plot shows the posterior proportion distribution, including the simulated proportion with a black dotted line.

To demonstrate `soibean`’s robustness, we downsampled the mixture of two bears to *∼*1.3*X, ∼*0.7*X* and *∼*0.25*X* coverage. `soibean` identifies the correct sources for every level of coverage (Supplementary Figure 17, 18 and 19) except for the 95% - 5% mixture at *∼* 0.25*X* coverage (see Supplementary Figure 19).

We repeated this experiment with a more dissimilar (83.4% similarity - Giant Panda bear and American Black bear) and a more similar (98.8% similarity - Ti-betan and Taiwan Black bear) mixture of two bears for all four coverage levels. Detailed results can be found in Supplementary Subsection 1.4 with Supplementary Figures 12 to 22. Generally, the higher the similarity, the higher the needed coverage for `soibean` to distinguish between sources successfully.

As a baseline, `HAYSTAC` performed comparably to `soibean` across all samples and mixtures, while `pathPhynder` was not developed to estimate more than one source. To demonstrate, we used the 55% - 45% mixture of the cave and the brown bear, where `pathPhynder` interprets the mixture as a single-source and identifies the mixture’s lowest common ancestor (see Supplementary Figure 23). `HAYSTAC`’s results for the two-source mixtures exhibited more variability at *∼*0.25*X* coverage, as outlined in Supplementary Table 1. This variability largely arises from a scarcity of uniquely mapped reads to species reference genomes at reduced coverage levels. In contrast, `soibean`’s pangenomic reference enhances its ability to navigate the challenges of unique identifiers by reducing reference bias, as discussed in Martiniano et al. [34].

We extended our testing for `soibean` to three-source and four-source samples. `soibean` shows continuous accuracy in identifying three sources and four sources with different levels of coverage. Detailed results are in the Supplementary Subsection 1.5.

### 2.2 Empirical Data

We demonstrate `soibean`’s efficacy on empirical data by showcasing its results on four published datasets. First, we reanalysed the 2-million-year-old sediment samples from Greenland’s Kap Københaven Formation [26]. The sample was sequenced using Illumina HiSeq and NovaSeq (ENA: PRJEB55522). The second dataset is an approximately 4000-year-old sediment sample from Qeqertasussuk, Greenland, sequenced with Illumina HiSeq (ENA: PRJEB13329) [55]. The third is a 25,000-year-old sediment sample from Satsurblia Cave in Georgia, sequenced using Illumina NovaSeq (ENA: PRJEB41420) [11]. Lastly, we reanalysed the metagenomic data sampled from pitch pieces used by a Mesolithic community in Huseby Klev, Sweden, dated to 9500 years ago, which was sequenced using Illumina Hiseq X (Bioproject: PRJNA994900) [25]. For each of the four samples, we downloaded the published data, trimmed adapters and merged the reads using `leeHom` [48], removed PCR duplicates and low-complexity reads using sga [56], and then inferred eukaryotic abundance using euka [65].

#### 2.2.1 Confirmatory Results

The first empirical sample describes different families of mammals, including the family Elephantidae, which was concluded to be a mastodon (*Mammut americanum*) using `pathPhynder` [33, 26]. Figure 4 A) shows `soibean`’s identification of the mastodon: first, the k-curve shows that a single source sufficiently explains the data; the likelihood does not show any significant increase for higher *k*s. Secondly, we show the estimated branch position for the mastodon. The higher the log-likelihood, the higher the confidence in the branch position. We visualise this confidence estimation in two ways: by colour (as seen in the legend) and by the position of dots (accepted MCMC moves) above or underneath the branch. If a dot is above the tree branch, the log-likelihood is higher than the median, and vice versa. If a point is positioned precisely on the branch, it equals the median log-likelihood of that MCMC chain.

**Figure 4:**
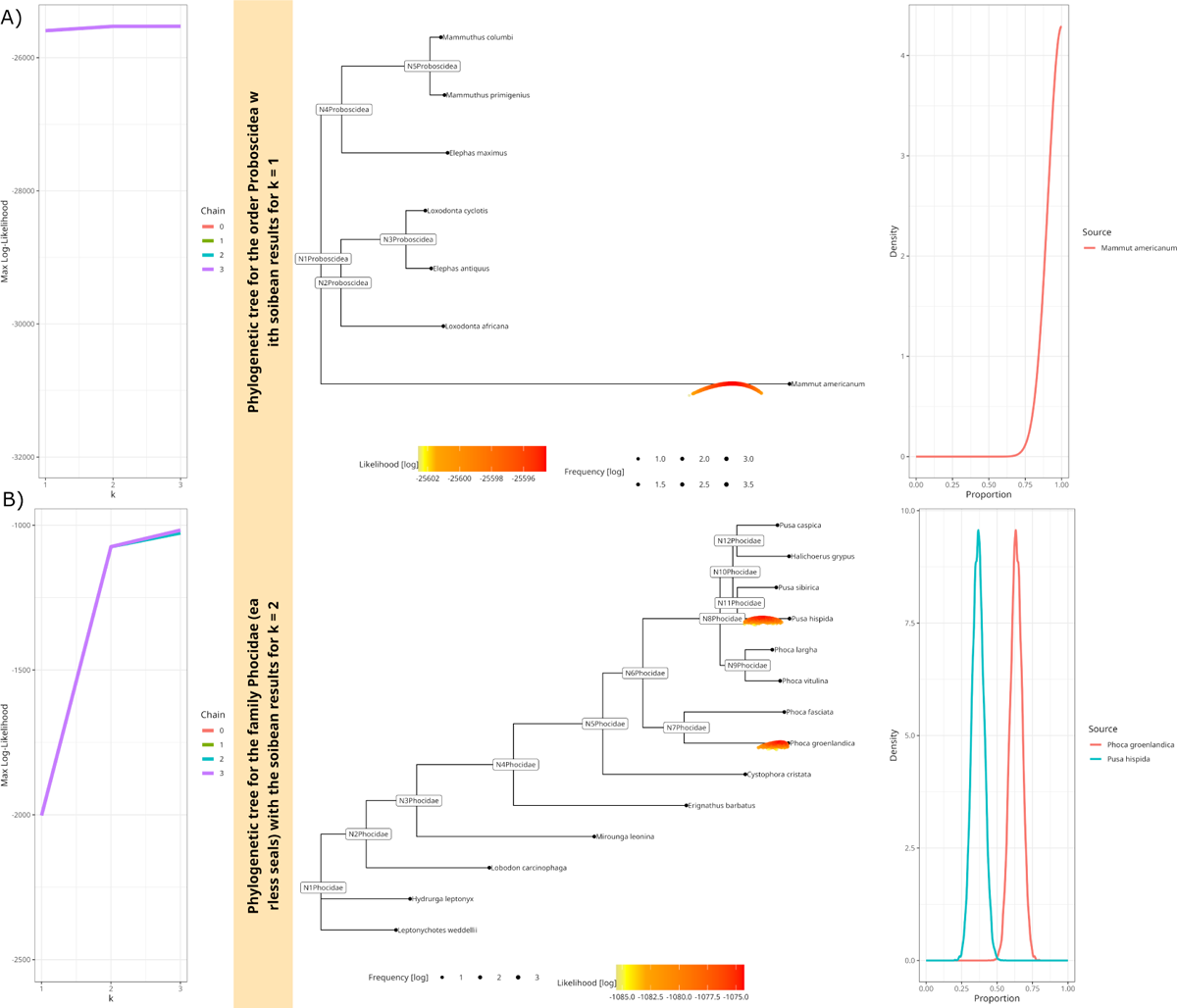
Confirmatory results for two independent studies A)soibean results for the 2-million-year-old empirical sample from Greenland. The filtered aDNA fragments from the order Proboscidea are analyzed using `soibean`’s standard parameters and a forced *k* = 3. The plotted k-curve shows there to be no more than one source present. The phylogenetic tree has every accepted move plotted along the most likely branch. A move plotted below the branch shows a lower likelihood than the median, and a move plotted above the branch has a higher likelihood. Moves are coloured by likelihood and show a clear result for the Mastodon, as seen in the original publication. We observe a slight divergence from the reference genome, which could be caused by the high levels of deamination observed in the sample or genetic divergence over time. B) `soibean`’s results for the 4000-year-old Greenlandic empirical sediment sample, where we analyzed the filtered aDNA fragments mapping to the family Phocidae. The k-curve on the left side of the plots clearly shows the presence of 2 contributing sources, which can be identified as the Harp seal *Phoca groenlandicus* and the Ringed seal *Pusa hispida*. This also aligns with the original publication. However, with `soibean`, we are able to add proportion estimates for the seals, which are found to be in a 60% −40% mixture.

The second sediment sample focused on evidence for the presence of bowhead whales. However, the original publication also identified different species of seals [55], which we focused on reanalysing. Figure 4 B) first shows the k-curve on the right side of the plot, which strongly suggests the presence of two sources in the sample but does not support a third source. When looking at the tree and the branch placement for the two sources, we can re-identify the Harp seal (*Phoca groenlandica*) and the Ringed seal (*Pusa hispida*), in a ratio of approximately 60% − 40%. The specific proportion often gets lost when mapping due to higher taxonomic classifications of reads. After duplicate removal, 166 aDNA fragments were mapped. The extremely low coverage does not allow us to define an exact position on the branch.

#### 2.2.2 Novel Results

For the third empirical cave sediment sample, the original publication focused on retrieving high-coverage mitochondrial genomes for three species (human, bison, and wolf) using shotgun metagenomics. They used phylogenetic placements to estimate the correct position in the species phylogenetic tree. Here, we focus on the reads that euka assigned to the family Bovidae. Supplementary Figure 28 shows a clear signature of two distinct Bovidae sources. `soibean` predicted the first one to be the European Bison (*Bison bonasus*), which is the same as found and analysed in the original publication. Additionally, `soibean` picked up on a small signal (5%) from the West Caucasian tur (*Capra caucasica*). The publication describes a signal from the genus *Ovis*. However, no conclusion was reached due to low coverage. Based on the mitochondrial data analysed with `soibean`, this second source could be the West Caucasian tur, which is native to Georgia and is believed to have been hunted at around the same time as the sample’s estimated age [43, 11]. We tried to add a secondary analysis to verify our results, where we concatenated the reference mitogenomes of bison (NCBI accession NC 014044.1) and tur (NCBI accession NC 020683.1), mapped all reads using SHRiMP [53] and extracted 54 reads mapping uniquely to the tur mitogenome. We plotted the deamination patterns for the alignment using bam2prof [50] (commands and parameters can be found in Supplementary Subsection 1.6). Due to the extreme sparsity of the data, the damage plot shows a high volume of noise from other substitutions (see Supplementary Figure 29). Subsequently, we cannot call a consensus from the tur data to place it phylogenetically for additional confirmation. This demonstrates the significance of `soibean` in identifying species from sparse data. The identification process opens up possibilities for employing laboratory enrichment methods to gain deeper insights into the ecological history of a sample.

The final reanalysed sample is a metagenomic sample from chewed pitch pieces of a Mesolithic community in Sweden. The original publication focused on the oral microbiome but also identified multiple eukaryotes, including foxes, salmon, deer, mallards, and apples, as potential food sources [25]. We used euka to reanalyse the dataset and detected all taxa of the original publication plus one additional taxon, Odontoceti (toothed whales). We extracted all aDNA fragments for this taxon and filtered for low-complexity and PCR duplicates. `soibean` estimated the filtered input to be a single-source sample from a harbour porpoise (*Phocoena phocoena*) (see Figure 5 A) k-curve, which shows a slight rise in the log-likelihood from *k* = 1 to *k* = 2, However, the estimated sources for *k* = 2 and *k* = 3 show geographically unlikely species: see Supplementary Figure 30 and 31), thus indicating that the data can be explained using a single-source. The harbour porpoise presently inhabits the Baltic Sea, and its bones have been found before at the same location (Huseby-Klev, Sweden) dated to the same period [16, 21]. We used the same input file to confirm our results and mapped it against the harbour porpoise reference genome (NCBI accession number: NC005280.1). We estimated the deamination patterns from the BAM file (see Figure 5 C) and Supplementary Figure 32) and afterwards used the alignment and damage profiles to estimate a damage-aware consensus sequence using endoCaller from schmutzi [51]. The consensus sequence was aligned with other reference genomes from the genus using prank and converted into a phylogenetic tree using RaxML (commands and parameters can be found in Supplementary Subsection 1.6). Figure 5 B) shows the source’s placement within the harbour porpoise clade. This finding suggests that the Mesolithic community in Sweden also used the harbour porpoise as a food source.

**Figure 5:**
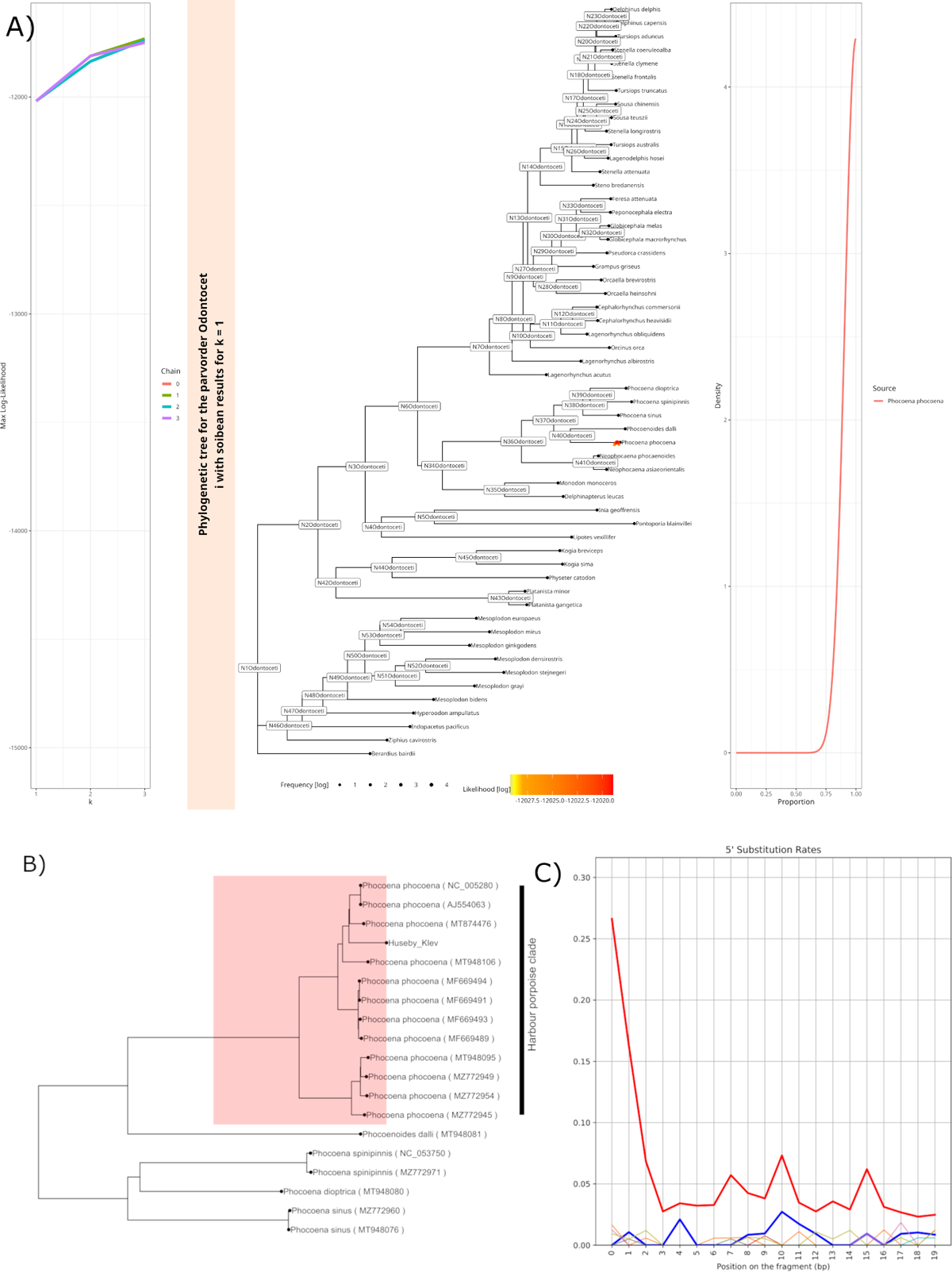
Novel results: A) `soibean` results for the 9500-year-old empirical metagenomic sample from pitch pieces found in Huseby-Klev on the northwestern coast of Sweden. We detected a previously unidentified taxon of toothed whales (Odontoceti). The filtered aDNA fragments from the parvorder Odontoceti are analyzed using `soibean`’s standard parameters and a forced *k* = 3. The plotted k-curve indicates a single-source sample. The phylogenetic tree has every accepted move plotted in colour and placement regarding their individual log-likelihood, displaying the most likely source to be the harbour porpoise (*Phocoena phocoena*). B) Maximum-likelihood phylogenetic tree plot with all available mitochondrial genomes for the porpoise including the consensus sequence for the Huseby-Klev sample. C) Deamination rates for the 5’end substitutions of the sample. The 3’end substitutions show a clearly elevated G to A pattern (see Supplementary Figure 32).

## 3 Discussion

`soibean` enhances aeDNA taxonomic specificity by distinguishing closely related species and estimating abundances with an MCMC algorithm, addressing the challenge of identification in low-abundance samples and advancing deep-time ecosystem insights. First, it is important to emphasise that our methodology, centred exclusively on mitochondrial aeDNA, may constrain the comprehensive analysis of exceedingly ancient samples [36]. In tests with simulated data, we confirmed `soibean`’s reliability for single-source data processing at minimum coverage depth of *∼*0.13*X*. This study aimed to determine the ancestral state in bears, a well-defined taxonomic group, ensuring high accuracy in ancestral state reconstructions. However, `soibean`’s performance may be less reliable with less-defined or more evolutionary divergent taxa, leading to uncertain ancestral sequence reconstructions. Caution is recommended when applying `soibean` to highly divergent or low-entropy taxa, often seen within the Arthropoda, due to the ongoing challenge of accurate arthropod classification in aeDNA research and lack of extensive testing in these areas.

In our study, we tested `soibean`’s ability to identify up to four sources within a genus, noting it theoretically could distinguish more but was limited by practical constraints. We faced challenges with closely related sources, as `soibean` needed higher coverage for accurate differentiation in taxa with low divergence due to few unique signature nodes. Improving the MCMC algorithm’s solution accuracy involved using signature node estimation for initialisation and increasing iterations, particularly vital when identifying over two sources and at lower coverages.

We recognise `soibean`’s higher computational demand and slower speed compared to other tools. However, its precision and reliability, especially for low-coverage Tetrapod and Arthropod aeDNA/aDNA samples, establish it as a crucial classification tool. Our visualisation scripts and diagnostic outputs aid in efficient result interpretation. Given the challenges in species classification due to aeDNA’s peculiarities, accurately quantifying uncertainty is vital. `soibean`’s use of Bayesian inference provides confidence levels for parameter estimation, proofing highly effective for aeDNA’s complex scenarios.

## 4 Methods

### 4.1 Database and Tree Construction

The database for `soibean` uses the same taxa introduced in the database for euka [65], which comprise 335 different taxa of tetrapodic and arthropodic eukaryotes (the taxon Galloanserae has been resorted since euka’s latest release). We used the same reference genomes as input to construct a multiple sequence alignment using PRANK version v.170427 [31], from which we constructed a phylogenetic tree using RAxML version 8.2.12 with the Hasegawa Kishino Yano (HKY) substitution model [17] (--HKY85) and estimated base frequencies [60]. The phylogenetic tree is used to infer the ancestral states for each taxon using FastML version 3.11 with the -mr flag for mitochondrial genomes and the -mh flag to use the HKY model [1]. We use the HKY substitution model for our guide tree as it allows for both nucleotide frequencies and the transition/transversion ratio to vary [17]. These features are crucial for the estimation of our guide tree and fitting our model because the transition/transversion ratio is much higher in mtDNA than in the autosomes [61] and because mtDNA is guanine-enriched on the heavy strand to aid in stabilizing RNA transcription [66, 29]. The HKY model allowed us to represent the variety of taxa used accurately. FastML outputs a multiple sequence alignment (MSA), including reconstructed ancestral states and the maximum-likelihood phylogenetic tree. We build a pangenome graph for each taxon from the new MSA with the added ancestral state sequences via the vg construct -M subcommand from the vg toolkit version 1.44.0 [10] (Supplementary Figure 1), resulting in 326 subgraphs (i.e. connected components), each subgraph corresponding to a different taxon for euka/`soibean`. A pangenome graph is defined as *G* = (*N, E, H*), where *N* are the nodes, *E* the edges, and *H* the graph’s embedded reference paths. Each node has a unique numerical identifier called node ID, which is internally assigned by vg. For a detailed overview of pangenome graphs applied to animal mitogenomes, see the Supplementary Material from Vogel et al. [65]. Each path in the graph corresponds to a mitogenome in the MSA and, therefore, a node in the phylogenetic tree. Inferred ancestral genomes correspond to internal nodes, whereas the mitogenomes used as input to the MSA are leaves in the phylogenetic tree. We merged connected components for each pangenomic graph into a single set of multiple independent graphs using the vg ids subcommand [10]. This resulted in a combined graph with *|N|* = 6889846, *|E|* = 10880449 and *|H|* = 10992.

We updated each taxon’s start and end node ID for our graph index and additionally empirically computed the base frequencies for each taxon to be used for our HKY-based likelihood model. A complete overview of the database construction and all scripts used can be found https://github.com/nicolaavogel/soibeanDatabase.

### 4.2 General Workflow

To describe `soibean`’s workflow, we will 1) specify the input data, 2) define how to access the database, and 3) describe the main function. 1) `soibean` accepts FASTQ input (single-end, paired-end interleaved, or paired-end separate) where the sequencing adapters have been removed, overlapping portions merged (see recommendations in [30]) and PCR duplicates removed. The FASTQ input should be DNA fragments previously assigned to the same higher taxonomic rank, e.g. all reads mapping to the taxon Ursidae (bears). This pre-filter is important as `soibean` does not have a model of spurious mappings due to bacterial DNA or DNA originating from another taxon. We recommend using euka or an alignment-based mapping + LCA approach to do the initial classification. 2) Once the taxon of interest is defined, the user can extract the pangenomic subgraph of the taxon of interest from the larger database using a provided bash script that takes the taxon name as input. The subgraph corresponding to this taxon is extracted from the combined pangenome graph by its start and end node IDs. The script automatically produces all index files required by vg giraffe [57]. Afterwards, the extracted taxon can be specified with the --dbprefix flag when using `soibean`. The specification of the database prefix simultaneously accesses the correct corresponding phylogenetic tree. 3) Once the input FASTQ mapping to the pangenomic component is done, we use a Markov Chain Monte Carlo (MCMC) sampling algorithm to estimate the most likely number of distinct contributing sources present in the sample and their respective placements on the (fixed) phylogenetic tree which is described in the subsections below. Test data can be found at https://github.com/nicolaavogel/soibeanDatabase and examples of the usage for `soibean` is provided on GitHub https://github.com/grenaud/vgan. During the development of `soibean`, we utilised the help of ChatGPT-4, a language model developed by OpenAI, for coding and debugging tasks. We carefully evaluated all proposed code before integration into the final software.

### 4.3 soibean Likelihood Function

Our model uses a maximum-likelihood framework to estimate the most likely number (denoted *k*) of contributing sources, their proportion ***θ*** (a length-k vector summing to one), and their most likely placements on branches ***β*** (a length-k vector of locations on the reference phylogenetic tree). Placements can be any position on the branches of the tree - not only nodes. This defines our model as *M* = (***β***, ***θ***). For instance, if we have two equally contributing sources, then *k* = 2 and ***θ*** = (0.5, 0.5). The phylogenetic placement ***β*** = (*β*_1_*, β*_2_) would then represent the placements on branches of the tree for each of the two sources.

We here use a uniform prior over the phylogenetic placement and abundance vectors. Both the prior and marginal probability are, therefore, constants, and according to Bayes’ Theorem, the posterior probability over the model parameters is consequently proportional to the likelihood (the probability of the data given the model parameters):

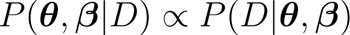

where the data, *D*, is the set of aligned reads. Briefly, we seek ***θ***, ***β*** that maximize *P* (*D|****θ***, ***β***). As we do not know the provenance of each read, we marginalise over each read for every source:

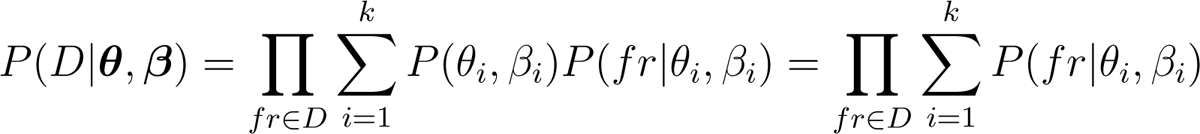

Specifically, a fragment *fr* can come from one of the *k* possible sources. The prior probability that a fragment is from source *i* and having placement *β_i_* is *P* (*θ_i_, β_i_*). The probability that a fragment *fr* has a specific sequence is *P* (*fr|β_i_, θ_i_*). Since we don’t know which of the *k* sources any given fragment is from, we compute the likelihood of a fragment by summing over these *k* possibilities. The overall likelihood for all fragments is then computed by multiplying their individual likelihoods, thus treating each DNA fragment as an independent observation and assuming duplicate fragments have been removed. The prior *P* (*θ_i_, β_i_*) can be omitted from the calculation because we use a flat (and hence constant) prior on both these parameters.

To calculate the likelihood of a branch placement on the tree, we make the simplifying assumption of treating each base *b* of fragment *fr* as independent, allowing us to multiply the probabilities of each nucleotide observation for each fragment:

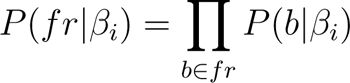

To compute the probability of a given nucleotide observation, we calculate the probability of observing the base given the placement. Any placement on the branch of a tree can viewed as a position between 2 nodes (not to be confused with nodes in the pangenome graph, which represent sequences), a derived node *N_D_*, which is closer to the leaves of the tree and an ancestral node *N_A_* which is closer to the root of the tree. Each tree node has a single reference path in the graph. For the nodes in the pangenomic graph, there are a certain number of reference paths that go through them. For our base *b*, there are two possibilities for any given tree node 1) the base was aligned to graph nodes associated with the tree node or 2) the base missed the alignment to all or certain graph nodes associated with the tree node. We discuss both cases.

#### *b* is missing certain graph nodes for a given reference path

Due to the nature of the pangenome graph structure, it may be the case that the base in question is on a node untraversed by the putative reference path. We term such nodes ‘unsupported’ by the path. In these cases, we treat 6/7 of all bases as a sequencing error with a probability of ɛ/3 and 1/7 of all bases as match with probability 1 *−ɛ*. *ɛ* is directly derived from the base quality reported from the sequencer. This is a slight update of the model for unsupported bases, which was used in HaploCart, another `vgan` subcommand [52], which we find to be more accurate empirically as the taxa used in the `soibean` database have a higher genetic divergence than human mitochondrial haplotypes.

#### *b* aligns to graph nodes for a given reference path

If the base *b* is aligned to a node associated with a path that corresponds to either tree node along the tree branch (namely either *N_D_*or *N_A_*), we compute the probability of either a match, a mismatch, a deletion, an insertion, an unresolved base or a softclip (an unaligned portion flanking an aligned fragment). For an aligned base, deletions and insertions have a probability of 0.02 based on an empirical study of human mitogenomes [28]. Unresolved bases, as well as softclips, are treated as sequencing errors with a probability of 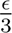.

We are left estimating the probability of an aligned base *b* being a match or a mismatch. Three events could change a nucleotide: a mutation occurring with a probability *µ*, ancient damage with a probability *δ*, or a sequencing error occurring with probability *ɛ*. A match would be the absence of all these events. However, it is also possible, but less likely, that a match occurred due to a mutation followed by damage, which reverted the base to the original one. We compute the probability of all these scenarios. This means we want to compute the probability of *P* (*b|b_g_*) where *b_g_* is the reference base.

We first look at the probability of a mutation given a position on our tree branch *t* under an HKY model [17]. A detailed explanation of how we calculate *µ* and the resulting probabilities for a match, transition or transversion can be found in Supplementary Subsection 2.1. The principal calculation is as follows: the further we move from *N_D_* towards its ancestor *N_A_*(the higher the value for *t*), the higher the probability of a mismatch caused by mutation. This follows the evolutionary model for a given taxon, allowing us to represent diverse and conserved taxa equally well with one algorithm. If a taxon has higher mitodiversity and longer branch lengths, the model is more lenient towards substitutions. Conversely, substitutions incur a lower likelihood of a taxon’s mitogenome being highly conserved. After considering the probability of a mutation, we denote *P* (*b_s_|b_g_*), where *b_s_* is our graph base after marginalising over every possibility of a mutation.

Following mutation, a mismatch can be explained by a deaminated base (*C→ U*, read by the sequencer as T, or *G→ A*) in the fragment. The probability of observing a mismatch explained by a deamination event is given by:

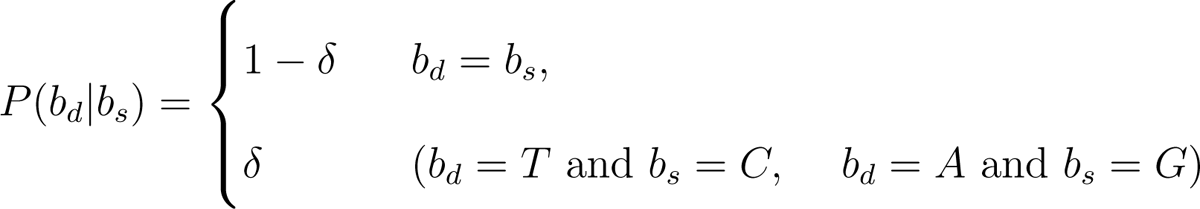

where *b_d_* is our graph base after the marginalisation of all possible cases of damage. *δ* depends on the base position within the fragment. The probability of a deamination event is higher at the 5’ end of the fragment for *C → T* substitutions and the 3’ end for *G → A* substitutions. We allow the user to provide damage rate matrices for their data to reflect the level of damage in their sample. The probabilities of deamination and sequencing error are independent of the tree placement. This allows us to precompute them for every alignment in the data at runtime. Again, the probability of observing either of the four bases following deamination is computed.

Following mutations and damage, we compute the probability of a sequencing error *ɛ* is derived from the base quality reported by the sequencer and denoted by:

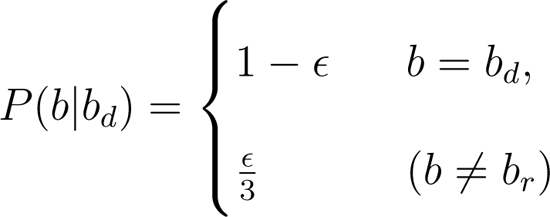

A marginalisation over each of the 4 bases following a potential sequencing error is performed to obtain our likelihood model and the probability of *b*.

Finally, we calculate the probability of the base for these two possibilities: (1) the source is *N_A_*, (2) the source is *N_D_*. The length of the branch from *N_A_* to *N_D_*is *t*, and the relative branch placement is *β_i_*, where 0 *≤ β_i_ ≤* 1. A value of *β_i_*= 1 would imply that we believe that the source was equal to *N_D_*, while *β_i_* = 0 would mean that the source was *N_A_*. The distance from the source to *N_D_* is *t_D_* = (1 *− β_i_*)*t*, while the distance from the source to *N_A_* is *t_A_* = *β_i_t*. We compute the product of *P* (*b|N_A_, t_A_*) and *P* (*b|N_D_, t_D_*) and calculate their weighted average across the entire aligned DNA fragment *r* consisting of *j* aligned bases denoted *b_i_*:

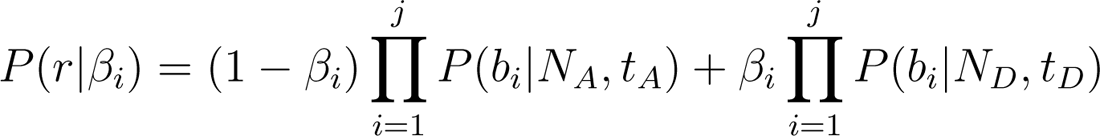

The product of all reads gives us the final likelihood of a single source. The case for multiple sources is found on page 20.

### 4.4 Signature Node Detection

`soibean` first maps the input FASTQ file to the subgraph corresponding to the taxon of interest. We count the number of aDNA fragments that align to the different signature node sets in the pangenome graph. We use the term ‘signature nodes’ in analogy with the concept of ‘signature genes’ in metagenomics to denote nodes in the graph only supported by one unique reference path. A signature node represents one or multiple bases in a position of the mitochondrial genome, which is unique to one species (reference genome) in the graph. We term the set of all signature nodes for a given reference sequence the ‘signature node set’.

Based on the total number of aligned aDNA fragments to a signature node set in the pangenome graph, we can estimate an initial number of distinct sources present in the sample. A signature node set must have a total frequency of more than 1% of the total alignments to the entire subgraph. This is implemented to reduce signature node predictions from noisy data. We set our first estimate of *k* as our maximum *k* value and run our MCMC sampling algorithm for every whole number from 0 to *k*. For a visual representation of the workflow, see Figure 1.

### 4.5 MCMC Sampling

The MCMC process is run *k* times, corresponding to the number of significant signature node paths identified (or alternatively, the number specified by the user if one is provided). For each MCMC run, four chains are deployed: the first chain is initialized at the identified signature nodes, whereas the subsequent three incorporate random nodes from the tree as starting points. The initial run of the MCMC (*k* = 1) initializes placement from the most prevalent signature node path. Subsequent runs incrementally include the next most frequent paths as starting points, continuing this pattern until *k* equals the total count of significant signature node paths. The initial proportion of each source is set at 1*/k*, and the initial branch position is sampled uniformly from [0, 1], where 0 signifies the ancestral state, and 1 is the derived state of the branch.

We use a Metropolis-Hastings algorithm [37, 18] for our MCMC sampling. Throughout the MCMC runs, new branch placements and source proportions are sampled.

The branches are normalised to unit length *t*, with the *N_A_* marked as 0 and the *N_D_* as 1. Proposed distances *d* to new placements ***β****^∗^* are drawn from a normal distribution, with a mean of 0 and the standard deviation linearly decreasing from 3 to 0.1 throughout the burn-in phase, then from 0.1 to 1*e^−^*^10^ in subsequent iterations. Should a proposed distance be negative, the transition is towards the root; conversely, a positive distance indicates a movement towards the leaves.

If the sum of our current position on the tree branch and the newly sampled distance is smaller than one (*d* + *β_i_ <* 1) or the difference between the current position and the sampled distance is higher than 0 (*d − β_i_ >* 0), we update our current position on the branch to the updated position on the branch (*β_i±d_*). If the sum of our current position on the tree branch and the newly sampled distance is larger than 1 (*d* + *β_i_ >* 1), we move to either of the two child branches of *N_D_* equiprobably.

In case *N_D_* is a leaf node, we sample a new move. When the difference between the current position on the tree branch and the sampled distance is negative (*d−β_i_ >* 0) we move to either the parent branch or the sibling branch of *N_A_* equiprobably. If *N_A_* is the root node, we sample a new move (see Figure 1).

If *k ̸*= 1, we simultaneously sample a new proportion vector ***θ^∗^***. Every proportion *θ^∗^* is sampled from *N* (*θ_i_,* 0.1) and *θ^∗^ ∈* [0, 1]. We use the sum of ***θ^∗^*** to normalise each *θ^∗^* so that ***θ^∗^*** = 1.

We calculate the new likelihood for *P* (*D|****β****^∗^,* ***θ****^∗^*). If *P* (*D|****β****^∗^,* ***θ****^∗^*) *> P* (*D|****β***, ***θ***) we accept the move, append the new parameter values to the sample file, and continue to the next iteration. If *P* (*D|****β****^∗^,* ***θ****^∗^*) *< P* (*D|****β***, ***θ***) we accept the move with probability 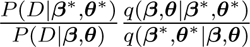. Here *q* is the proposal distribution for moves in parameter space, and *q*(*X|Y*) is the probability of proposing *X* when the current parameter values are *Y*. If this distribution is symmetric, i.e. *q*(*X|Y*) = *q*(*Y |X*), the q-terms cancel. In the Supplementary Material Section 2.2, we show that our proposal sampling scheme is indeed symmetric, so this factor is exactly one and can, therefore, be disregarded.

By default, we perform 500,000 MCMC iterations and discard the first 75,000 iterations as burn-in. These values can be overridden by the user via command-line options. Notably, lower values of *k* should require a lower number of iterations for convergence.

#### 4.4.1 MCMC Diagnostics

After inference is complete, it is important to assess whether the chains managed to converge to the true underlying posterior distribution. This can be assessed individually for each parameter of the model. To assess the convergence of phylogenetic placement, we keep track of the patristic distance from the accepted position to all the leaves on the tree (the patristic distance is the distance along the tree between two nodes, i.e., the sum of lengths of the branches on the path between those nodes [9]). Specifically, we compute a vector where the elements are the patristic distances from the accepted position to each of the leaf nodes. We also compute a vector giving the patristic distances from the *root node* to each of the leaves. This is done for every one hundred iterations after the burn-in period. We then compute the Euclidean distance between these two vectors of patristic distances. If a trace plot of this Euclidean distance against MCMC iteration shows any upward or downward trend, then that would indicate a lack of convergence. If phylogenetic placement *has* converged then the trace plot should move around a constant value. We also provide the user with the chain statistics, including the mean, median, 95% credible intervals, variance, autocorrelation, and effective sample size (ESS) for each chain and each estimated parameter. We warn the user if the ESS for a chain is below 200 [35], as it could mean the estimates are insufficiently precise.

Another important and widely-used metric for assessing convergence is called *R*^^^. We compute *R*^^^ for both ***θ*** and ***β*** across chains. The measure essentially compares the within-chain variance with the between-chain variance; a value close to 1 indicates convergence [12, 5]. Values larger than 1.05 [62] may indicate non-converging chains.

## 5 Declarations

### 5.1 Software Versions

We used PRANK version v.170427, RAxML version 8.2.12, and FastML version 3.11. Our vg version was 1.44.0 - ‘Solara’, SPIMAP version 1.2 [47] and `vgan` version 3.0.0 - Fagiolo. We used `HAYSTAC` version v0.4.8 and `pathPhynder` version 1a with BWA version 0.7.17. Our simulated data was created using `gargammel` version 1.1.2, ART version 2.5.8, and leeHom version 1.2.15. Additional analysis was done using SHRiMP version 2.2.2., bam2prof version 1.5.4 and schmutzi’s endoCaller version 1.5.6. All plots were produced using R version 4.3.1 - ‘Beagle Scouts’.

### 5.2 Data Availability

`vgan` can be built from source or downloaded as a static binary from https://github.com/grenaud/vgan as well as Zenodo https://doi.org/10.5281/zenodo. 7875929 [24]. It is also available on BioConda https://bioconda.github.io/recipes/vgan/README.html and as a Docker image. Database construction scripts, as well as all simulated data, are available at https://github.com/nicolaavogel/soibeanDatabase or from Zenodo https://zenodo.org/records/10828227 [64].

## Acknowledgements

We would like to acknowledge that this research and the PhD scholarships of Nicola Alexandra Vogel and Joshua Daniel Rubin were funded by the Novo Nordisk Data Science Investigator grant number NNF20OC0062491. We thank the Technical University of Denmark’s Department of Health Technology for additional funding. Additionally, we would like to thank Rui Martiniano and Bianca De Sanctis for their thoughtful comments on our manuscript.

## 5.3 Authors’ Contributions

N.A.V., J.D.R., and G.R. developed and implemented the method. M.W.P. and A.G.P. supported implementation and data interpretation. N.A.V. conducted all tests. P.W.S. provided IT infrastructure support. All authors approved the final manuscript.

