## Supplementary Material for "soibean: High-resolution Taxonomic Identification of Ancient Environmental DNA Using Mitochondrial Pangenome Graphs"

### 1 Supplementary Results

#### 1.1 Computational Performance

`soibean`'s computational performance highly depends on the used reference graph, the size of the input FASTQ and the number of iterations specified for the MCMC. Supplementary Figure 27 shows a comparison of user time in hours for three different sizes of FASTQ input and three different numbers of iterations set for the MCMC. Each run multithreaded to 20 cores. `soibean` assesses the log-likelihood for the FASTQ-input in each iteration of the MCMC to take a new placement on the branch into account. Given our HKY model (see Method Section 2.1, our model needs to recompute the log-likelihood for every base of every fragment associated with the source. This is time-intensive; however, it provides us with extremely accurate estimations. Additionally, `soibean` can be multi-threaded, improving time performance.

We have observed that `soibean`'s memory consumption depends primarily on the reference graph used. On average, `soibean` uses about 1.5Gb of memory. This varies depending on the size of the input graph.

#### 1.2 Benchmark

We benchmarked against `pathPhynder` with standard parameters [6] for our single-source simulations. We followed the tool's recommended guidelines and the approach taken in Kjær et al. [3]. We used `BWA ALN` [5] for mapping to the consensus sequence. The `pathPhynder` best path results for our single-source samples are visualised in Supplementary Figures 2, 3, 4, 5, and 6. `pathPhynder`'s maximum likelihood results for the single-source samples are visualised in Supplementary Figures 7, 8, 9, 10, and 11.

To the best of our knowledge, no existing tool estimates more than one source on a phylogenetic tree. Therefore, we used `HAYSTAC` as a baseline model. The tool specialises in classifying highly similar samples [1]. We used `HAYSTAC`'s standard parameters and recommended thresholds for their coverage evenness filter; additionally, we specified that a source needs at least five uniquely mapped reads to be considered present. All results are summarised in Supplementary Table 1.

#### 1.3 Benchmarking Commands

##### 1.3.1 soibean

```
./make_graph_files.sh [database name]

vgan soibean -fq1 [simulation.fq.gz] -t 20 --dbprefix [database name]
-o [output name]
```

##### 1.3.2 HAYSTAC

```
haystac sample --output [sample_dir] --fastq [simulation.fq.gz]
--core 20 --trim-adapter False

haystac analyse --mode abundances --database [database]
--sample [sample_dir] --output [ouput_dir] --bowtie2-threads 20
--aDNA --cores 20
```

##### 1.3.3 pathPhynder - best path method

```
bwa aln -l 1024 -n 0.001 -t 10 [consensus.fasta] [simulation.fq.gz]
| bwa samse [consensus.fasta] - [simulation.fq.gz] | samtools view
-F 4 -q 25 -@ 10 -uS - | samtools sort -@ 10 -o [output.sort.bam]
```

```
phynder -B -o[output.snp] [tree.newick] [input.vcf]
```

```
pathPhynder -s prepare -i [tree.newick] -p [taxa_pathphynder_tree]
-f [output.snp] -r [consensus.fasta]
```

```
pathPhynder -s all -t 10 -i [tree.newick]
-p [taxa_pathphynder_tree] -b [output.sort.bam] -r [consensus.fasta]
```

###### 1.3.4 pathPhynder - maximum likelihood method

```
bwa aln -l 1024 -n 0.001 -t 10 [consensus.fasta] [simulation.fq.gz]
| bwa samse [consensus.fasta] - [simulation.fq.gz] | samtools view
-F 4 -q 25 -@ 10 -uS - | samtools sort -@ 10 -o [output.sort.bam]
```

```
phynder -B -o[output.snp] [tree.newick] [input.vcf]
```

```
pathPhynder -s prepare -i [tree.newick] -p [taxa_pathphynder_tree]
-f [output.snp]
```

```
Rscript R/make_vcf.R intree_folder2/ 1 [sample.vcf]
```

```
./phynder/phynder -q [output.vcf] -p 0.01 -o [query.output]
[tree.newick] [input.vcf]
```

#### 1.4 Two source simulation

Additionally, we repeated this experiment with a distantly related mix of a Giant Panda bear and an American Black bear (83.4% similarity), as well as a very closely related mix of the Tibetan and Taiwan Black bear (98.8% similarity). Both mixtures were simulated in the same four proportions: $95\% - 5\%, 85\% - 15\%, 75\% - 25\%$ and $55\% - 45\%$ with an average coverage of $\sim 2.5X$ (1000 aDNA fragments) as in Pedersen et al. [7]. At this coverage, we obtain correct identification for the Giant Panda and American Black bear mixture at any proportion (Supplementary Figure 12), while we misidentify the closely related sample for the $95\% - 5\%$ and the $85\% - 15\%$ mixtures of the Asian Black bears (Supplementary Figure 13).

We repeat `soibean`'s robustness test by downsampling both mixtures to $\sim 1.3X$, $\sim 0.7X$ and $\sim 0.25X$ coverage. `soibean` can identify the mixture of the Giant Panda and the American Black bear for every mixture percentage until the lowest coverage (see Supplementary Figures 14, 15,16). For the most similar sample of Asian Black bears, we can observe that the $95\% - 5\%$ can never be identified, while the rest can be placed on the correct tree branch (see Supplementary Figures 20, 21 and 22), when downsampling.

#### 1.5 Three and Four Source Simulations

We simulated a three-source sample from a family of moths (Saturniidae), including the two emperor moths *Gonimbrasia tyrrhea* and *Gonimbrasia belina*. Additionally, we added the ancestral state N7 to the mixture and simulated a total of 1500 aDNA fragments, averaging a coverage of $\sim 4X$. Mixture proportions were $47\% - 33\% - 20\%$. For our simulations of a four-source sample, we used the family of seals (Phocidae); specifically, we sampled 500 aDNA fragments for each of the four earless seals, namely *Phoca largha*, *Phoca vitulina*, *Phoca groenlandica* and *Pusa hispida* creating a mixture of 25% each and a total coverage of about $\sim 5.4X$. These numbers of simulated fragments were up-scaled from Pedersen et al. [7] as we could not find an empirical aeDNA study identifying three or more sources from one family.

Supplementary Figure 24 shows `soibean`'s results for three (top) and four (bottom) simulated sources, clearly identifying the correct placements and proportions for each. A warning is produced in case a chain's effective sample size (ESS) is below 200. This warning was triggered for the four-source sample. The ESS is essentially the number of independent MCMC samples (accounting for autocorrelation). A low ESS means that the quantiles of the posterior distribution will be poorly estimated (especially quantiles in the tails of the posterior, such as the 5% and 95%) and is an indication that the MCMC should be run for more iterations.

Again, we downsampled both samples to demonstrate `soibean`'s robustness. The three-source samples can be consistently identified until a coverage of $\sim 1X$. At lower coverage, it becomes more difficult to identify the branch placement for the ancestral state N7 and more branch positions on the surrounding nodes of N7 are accepted (Supplementary Figure 25). For the four-source samples, `soibean` can identify the correct sources for each downsampled coverage if the signature node prediction is used to initiate the starting position on the tree (see Supplementary Figures 26). However, if we initialise randomly, `soibean` does not converge to the correct branch placements. Our diagnostics clearly identify the highest log-likelihood with the correct branch placements in the phylogenetic tree, suggesting that a higher number of iterations is necessary to converge to the true underlying posterior.

Our baseline model identifies the two emperor moths for the three-source sample but does not pinpoint the ancestral state, as seen in the single-source sample (see Supplementary Table 1) for every simulated coverage. For the four-source sample, the baseline model identifies all four species down to coverage of $\sim 1.3X$.

#### 1.6 Emprical Data: Commands

##### 1.6.1 soibean

```
./make_graph_files.sh [database name]
```

```
vgan soibean -fq1 [input.fastq] --dbprefix [database name] -t 20
-o [output name] -k 3
```

##### 1.6.2 Phylogenetic Placement

```
SHRiMP_2_2_2/bin/gmapper -N 4 -o 1 --single-best-mapping --sam-unaligned
--fastq --sam --no-qv-check  --qv-offset 33  [input.fastq] [reference.fa]
| samtools view -bS  -F4 /dev/stdin | samtools fillmd -b  /dev/stdin
[reference.fa] | samtools sort /dev/stdin > [output.bam]
samtools index  [output.bam]

bam2prof -double  -5p [output_5p.prof] -3p [output_3p.prof] [output.bam]

endoCaller -seq [output.fa] -log [output.log] -name [name] -deam5p [output_5p.pro
-deam3p [output_3p.prof] [reference.fa] [output.bam]

cat [output.fa] [phylogeneticSeqs.fa] > [output_all.fa]

prank -d=[output_all.fa] -o=[output_all.prank] -showall -DNA

RAxML-8.2.12/raxmlHPC-PTHREADS -s [output_all.prank].best.fas --HKY85
-m GTRGAMMA -n [output_prefix] -w [output_dir] -p 76 -T 40 -d
```

### 2 Supplementary Methods

#### 2.1 HKY model

We are interested in finding the most likely placement of ancient species on a precomputed mitochondrial phylogenetic tree. To achieve this we use the HKY substitution model to compute the probability of an observed, ancient DNA sequence, arising by substitutions occurring along a branch of the tree, some distance from one of the existing nodes.

The instantaneous rate matrix for the HKY model can be written as [2, 8]:

$$\boldsymbol{Q} = \mu \begin{bmatrix} - & \pi_C & \kappa\pi_G & \pi_T \\ \pi_A & - & \pi_G & \kappa\pi_T \\ \kappa\pi_A & \pi_C & - & \pi_T \\ \pi_A & \kappa\pi_C & \pi_G & - \end{bmatrix}$$

The rows and columns of $\boldsymbol{Q}$ correspond to the nucleotides A, C, G, and T in that order. An element, $Q_{ij}$, gives the rate at which nucleotide $i$ changes to nucleotide $j$, and values are in units of substitutions per site per time unit. Elements on the diagonal (not shown here) are set such that all rows sum to zero: $Q_{ii} = -\sum_{j \neq i} Q_{ij}$. In the expressions above $\kappa$ is the ratio between the rates of transition-type and transversion-type substitutions, while $\pi_A, \pi_C, \pi_G$, and $\pi_T$ are the equilibrium nucleotide frequencies. The multiplicative factor $\mu$ will, in part, depend on the units of time used in the matrix (e.g., rates measured per century will be 100 times larger than rates measured per year).

Given the rate matrix $\boldsymbol{Q}$ it is possible to compute the transition probability matrix $\boldsymbol{P}(t)$ for a given branch length $t$: $\boldsymbol{P}(t) = e^{\boldsymbol{Q}t}$. In this expression, $t$ has to be expressed in the same time units as $\boldsymbol{Q}$. Since branch lengths in the pre-computed phylogeny are expressed as the expected change per site (and not in some unit of time), we first normalise $\boldsymbol{Q}$ correspondingly. Specifically, this is done by setting $\mu$ such that the expected rate of change, $\rho$, is 1: If $\rho = 1$ then the expected amount of change on a branch of length $t$ will be: $\nu = \rho t = t$. This is equivalent to having chosen a unit of time such that the expected change on a branch ($\nu$, measured in substitutions per site) is numerically identical to the branch length ($t$, measured in those time units).

The expected rate of change for the HKY model is [2]:

$$\begin{aligned}\rho &= \sum_i \pi_i \sum_{j \neq i} Q_{ij} \\ &= 2(\kappa\pi_A\pi_G + \kappa\pi_C\pi_T + \pi_A\pi_C + \pi_A\pi_T + \pi_C\pi_G + \pi_G\pi_T)\mu\end{aligned}$$

Setting this equal to 1, and isolating $\mu$, we find:

$$\mu = \frac{1}{2(\kappa\pi_A\pi_G + \kappa\pi_C\pi_T + \pi_A\pi_C + \pi_A\pi_T + \pi_G\pi_C + \pi_G\pi_T)}$$

Using $\boldsymbol{P}(t) = e^{\boldsymbol{Q}t}$ one can derive the following expressions for the probability of observing nucleotide $b$, in an ancient taxon, given the presence of the nucleotide $r$, at a distance of $t$, in a node on the tree:

$$\begin{cases} \pi_r + \pi_r(\frac{1}{\Pi_r} - 1)e^{-\mu t} + (\frac{\Pi_r - \pi_r}{\Pi_r})e^{\mu t A} & (b = r), \\ \pi_r + \pi_r(\frac{1}{\Pi_r} - 1)e^{-\mu t} - (\frac{\pi_r}{\Pi_r})e^{-\mu t A} & (b \neq r, \text{transition}), \\ \pi_r(1 - e^{-\mu t}) & (b \neq r, \text{transversion}) \end{cases}$$

Here $A = 1 + \Pi_r(\kappa - 1)$, with $\Pi_r = \pi_A + \pi_G$ (if $r$ is A or G) and $\Pi_r = \pi_C + \pi_T$ (if $r$ is C or T). The branch length $t$ is the distance between the ancient taxon and a nearby node on the tree and is expressed in units of expected change as explained above. Based on previous analyses of human mitochondrial evolution, we set $\kappa$ to 22 [4]. The equilibrium nucleotide frequencies are set to their empirical values in the data set at hand. The value of $\mu$ is computed from $\kappa$ and $(\pi_A, \pi_C, \pi_G, \pi_T)$ according to the expression above.

#### 2.2 Proof That the Proposal Sampling Distribution $q(\boldsymbol{\beta}^*, \boldsymbol{\theta}^* | \boldsymbol{\beta}, \boldsymbol{\theta})$ is Symmetric

We present a quick argument that the proposal sampling distribution is symmetric. In other words, we aim to argue that for Markov states $X$ and $Y$ in our parameter space, sampling $Y$ from state $X$ has a probability equal to that of sampling $X$ from state $Y$. To see this, recall that in our sampling scheme, we first fix a distance to move and a direction (ancestral or derived) in which to move that distance. Note also that for a given proposal move, we do not allow any self-intersecting paths on the tree. Therefore, given the bifurcating nature of tree topologies, there will always only be one possible way to sample $Y$ from $X$ for any Markov states $X, Y$. Along this path, there will be some number of bifurcations in which one of the two possible forks will be followed. The probability of taking any given fork is $\frac{1}{2}$. Since there are the same number of forks on the unique path from $Y$ to $X$ as there are from $X$ to $Y$, it clearly follows that the probability of sampling $X$ from $Y$ is identical to that of sampling $Y$ from $X$.

#### 3 Supplementary Table

| Reads | Average coverage | Proportion | Simulated Source(s) | Predicted Source(s) |
|---|---|---|---|---|
| 500 | 1.3X | 1 | Ancestral state *N4* | *U. americanus*<br>*M. ursinus*<br>*N13*<br>*U. spelaeus*<br>*U. arctos* |
| 250 | 0.66X | 1 | Ancestral state *N4* | *H. malayanus*<br>*U. americanus*<br>*M. ursinus* |
| 75 | 0.2X | 1 | Ancestral state *N4* | *U. americanus* |
| 50 | 0.13X | 1 | Ancestral state *N4* | None |
| 10 | 0.026X | 1 | Ancestral state *N4* | None |
| 1000 | 2.6X | 55-45 | *U. thibetanus formosanus*<br>*U. thibetanus thibetanus* | *U. thibetanus formosanus*<br>*U. thibetanus thibetanus* |
| 500 | 1.3X | 55-45 | *U. thibetanus formosanus*<br>*U. thibetanus thibetanus* | *U. thibetanus formosanus*<br>*U. thibetanus thibetanus* |
| 250 | 0.66X | 55-45 | *U. thibetanus formosanus*<br>*U. thibetanus thibetanus* | *U. thibetanus formosanus*<br>*U. thibetanus thibetanus* |

| | | | | |
|---|---|---|---|---|
| 100 | 0.26X | 55-45 | *U. thibetanus formosanus* *U. thibetanus thibetanus* | *U. thibetanus thibetanus* |
| 1000 | 2.6X | 55-45 | *U. spelaeus* *U. arctos* | *U. spelaeus* *U. arctos* |
| 500 | 1.3X | 55-45 | *U. spelaeus* *U. arctos* | *U. spelaeus* *U. arctos* |
| 250 | 0.66X | 55-45 | *U. spelaeus* *U. arctos* | *U. spelaeus* *U. arctos* |
| 100 | 0.26X | 55-45 | *U. spelaeus* *U. arctos* | *U. spelaeus* *U. arctos* |
| 1000 | 2.6X | 55-45 | *A. melanoleuca* *U. americanus* | *A. melanoleuca* *U. americanus* |
| 500 | 1.3X | 55-45 | *A. melanoleuca* *U. americanus* | *A. melanoleuca* *U. americanus* |
| 250 | 0.66X | 55-45 | *A. melanoleuca* *U. americanus* | *A. melanoleuca* *U. americanus* |
| 100 | 0.26X | 55-45 | *A. melanoleuca* *U. americanus* | *A. melanoleuca* *U. americanus* |

| | | | | |
|---|---|---|---|---|
| 1000 | 2.6X | 75-25 | *U. thibetanus formosanus* *U. thibetanus thibetanus* | *U. thibetanus formosanus* *U. thibetanus thibetanus* |
| 500 | 1.3X | 75-25 | *U. thibetanus formosanus* *U. thibetanus thibetanus* | *U. thibetanus formosanus* *U. thibetanus thibetanus* |
| 250 | 0.66X | 75-25 | *U. thibetanus formosanus* *U. thibetanus thibetanus* | *U. thibetanus formosanus* *U. thibetanus thibetanus* |
| 100 | 0.26X | 75-25 | *U. thibetanus formosanus* *U. thibetanus thibetanus* | *U. thibetanus formosanus* |
| 1000 | 2.6X | 75-25 | *U. spelaeus* *U. arctos* | *U. spelaeus* *U. arctos* |
| 500 | 1.3X | 75-25 | *U. spelaeus* *U. arctos* | *U. spelaeus* *U. arctos* |
| 250 | 0.66X | 75-25 | *U. spelaeus* *U. arctos* | *U. spelaeus* *U. arctos* |
| 100 | 0.26X | 75-25 | *U. spelaeus* *U. arctos* | *U. spelaeus* *U. arctos* |
| 1000 | 2.6X | 75-25 | *A. melanoleuca* *U. americanus* | *A. melanoleuca* *U. americanus* |

| | | | | |
|---|---|---|---|---|
| 500 | 1.3X | 75-25 | *A. melanoleuca U. americanus* | *A. melanoleuca U. americanus* |
| 250 | 0.66X | 75-25 | *A. melanoleuca U. americanus* | *A. melanoleuca U. americanus* |
| 100 | 0.26X | 75-25 | *A. melanoleuca U. americanus* | *A. melanoleuca U. americanus* |
| 1000 | 2.6X | 85-15 | *U. thibetanus formosanus U. thibetanus thibetanus* | *U. thibetanus formosanus U. thibetanus thibetanus* |
| 500 | 1.3X | 85-15 | *U. thibetanus formosanus U. thibetanus thibetanus* | *U. thibetanus formosanus U. thibetanus thibetanus* |
| 250 | 0.66X | 85-15 | *U. thibetanus formosanus U. thibetanus thibetanus* | *U. thibetanus formosanus* |
| 100 | 0.26X | 85-15 | *U. thibetanus formosanus U. thibetanus thibetanus* | *U. thibetanus formosanus* |

| | | | | |
|---|---|---|---|---|
| 1000 | 2.6X | 85-15 | *U. spelaeus* *U. arctos* | *U. spelaeus* *U. arctos* |
| 500 | 1.3X | 85-15 | *U. spelaeus* *U. arctos* | *U. spelaeus* *U. arctos* |
| 250 | 0.66X | 85-15 | *U. spelaeus* *U. arctos* | *U. spelaeus* |
| 100 | 0.26X | 85-15 | *U. spelaeus* *U. arctos* | *U. spelaeus* |
| 1000 | 2.6X | 85-15 | *A. melanoleuca* *U. americanus* | *A. melanoleuca* *U. americanus* |
| 500 | 1.3X | 85-15 | *A. melanoleuca* *U. americanus* | *A. melanoleuca* *U. americanus* |
| 250 | 0.66X | 85-15 | *A. melanoleuca* *U. americanus* | *A. melanoleuca* |
| 100 | 0.26X | 85-15 | *A. melanoleuca* *U. americanus* | *A. melanoleuca* *U. americanus* |

| | | | | |
|---|---|---|---|---|
| 1000 | 2.6X | 95-5 | *U. thibetanus formosanus* *U. thibetanus thibetanus* *M. ursinus* *A. melanoleuca* | *U. thibetanus formosanus* *U. thibetanus thibetanus* *M. ursinus* *A. melanoleuca* |
| 500 | 1.3X | 95-5 | *U. thibetanus formosanus* *U. thibetanus thibetanus* | *U. thibetanus formosanus* |
| 250 | 0.66X | 95-5 | *U. thibetanus formosanus* *U. thibetanus thibetanus* | *U. thibetanus formosanus* |
| 100 | 0.26X | 95-5 | *U. thibetanus formosanus* *U. thibetanus thibetanus* | *U. thibetanus formosanus* |
| 1000 | 2.6X | 95-5 | *U. spelaeus* *U. arctos* | *U. spelaeus* *U. arctos* |
| 500 | 1.3X | 95-5 | *U. spelaeus* *U. arctos* | *U. spelaeus* *U. arctos* |
| 250 | 0.66X | 95-5 | *U. spelaeus* *U. arctos* | *U. spelaeus* |
| 100 | 0.26X | 95-5 | *U. spelaeus* *U. arctos* | *U. spelaeus* |

| | | | | |
|---|---|---|---|---|
| 1000 | 2.6X | 95-5 | *A. melanoleuca U. americanus* | *A. melanoleuca U. americanus* |
| 500 | 1.3X | 95-5 | *A. melanoleuca U. americanus* | *A. melanoleuca U. americanus* |
| 250 | 0.66X | 95-5 | *A. melanoleuca U. americanus* | *A. melanoleuca U. americanus* |
| 100 | 0.26X | 95-5 | *A. melanoleuca U. americanus* | *U. americanus* |
| 1500 | 4X | 47-33-20 | *G. tyrrhea G. belina N7* | *G. tyrrhea G. belina G. maja B. alcinoe G. cytherea N. wahlbergi* |
| 750 | 2X | 47-33-20 | *G. tyrrhea G. belina N7* | *G. tyrrhea G. belina B. alcinoe G. maja G. cytherea* |

| | | | | |
|---|---|---|---|---|
| 375 | 1X | 47-33-20 | *G. tyrrhea*<br>*G. belina*<br>*N7* | *G. tyrrhea*<br>*G. belina*<br>*B. alcinoe* |
| 150 | 0.4X | 47-33-20 | *G. tyrrhea*<br>*G. belina*<br>*N7* | *G. tyrrhea*<br>*G. belina* |
| 2000 | 5.2X | 25-25-25-25 | *P. fasciata*<br>*P. groen-landica*<br>*P. vitulina*<br>*P. largha* | *P. fasciata*<br>*P. groen-landica*<br>*P. vitulina*<br>*P. largha* |
| 1000 | 2.6X | 25-25-25-25 | *P. fasciata*<br>*P. groen-landica*<br>*P. vitulina*<br>*P. largha* | *P. fasciata*<br>*P. groen-landica*<br>*P. vitulina*<br>*P. largha* |
| 500 | 1.3X | 25-25-25-25 | *P. fasciata*<br>*P. groen-landica*<br>*P. vitulina*<br>*P. largha* | *P. fasciata*<br>*P. groen-landica*<br>*P. vitulina*<br>*P. largha* |
| 200 | 0.55X | 25-25-25-25 | *P. fasciata*<br>*P. groen-landica*<br>*P. vitulina*<br>*P. largha* | *P. fasciata*<br>*P. groen-landica*<br>*P. vitulina* |

Table 1: Results table for the baseline model `HAYSTAC` for every simulated test.

### 4 Supplementary Figures

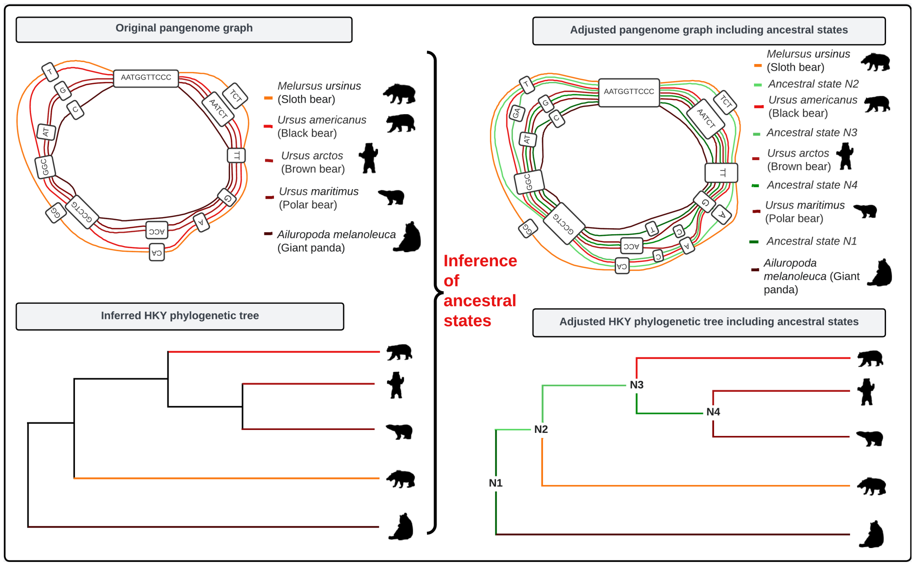


Figure 1: Workflow of a pangenome graph including ancestral states. The figure shows the original pangenome graph, including their phylogenetic tree, and follows with an adjusted pangenome graph, where ancestral state sequences are included and the corresponding phylogenetic tree.

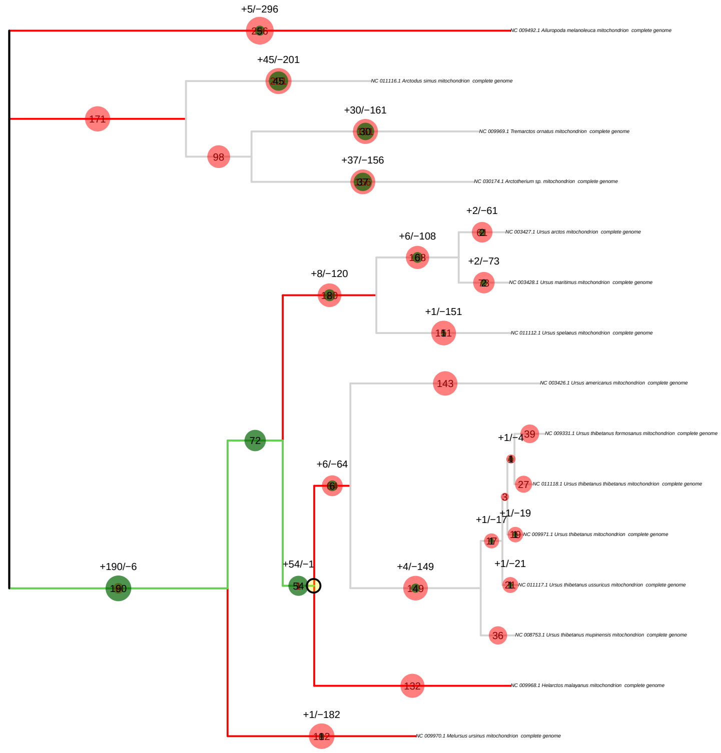


Figure 2: `pathPhynder` best path method results for the simulated ancient single-source sample N4 at $\sim 1.3x$ coverage. `pathPhynder` shows the best path for the given sample, which ends at the correct ancestral node N4.

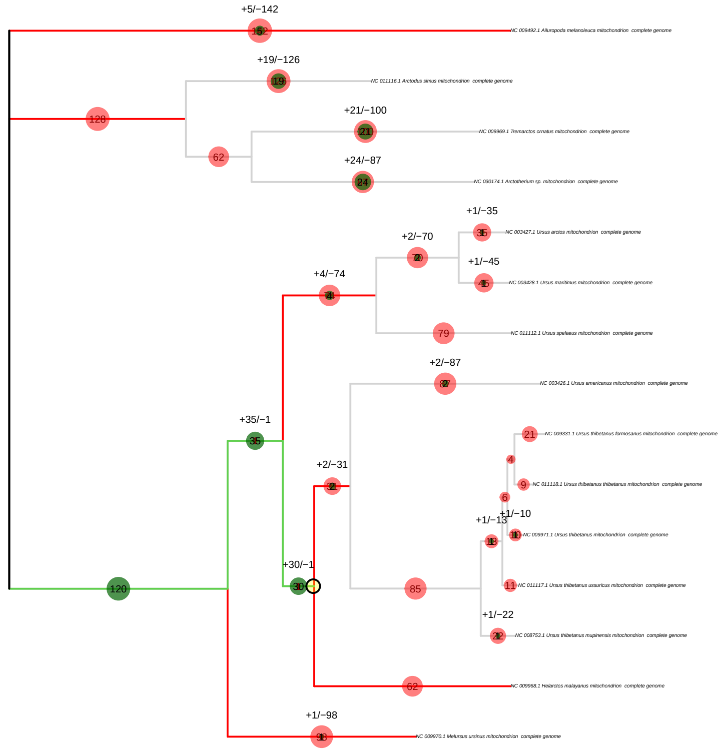


Figure 3: `pathPhynder` best path method results for the simulated ancient single-source sample N4 at $\sim 0.66X$ coverage. `pathPhynder` shows the best path for the given sample, which ends at the correct ancestral node N4.

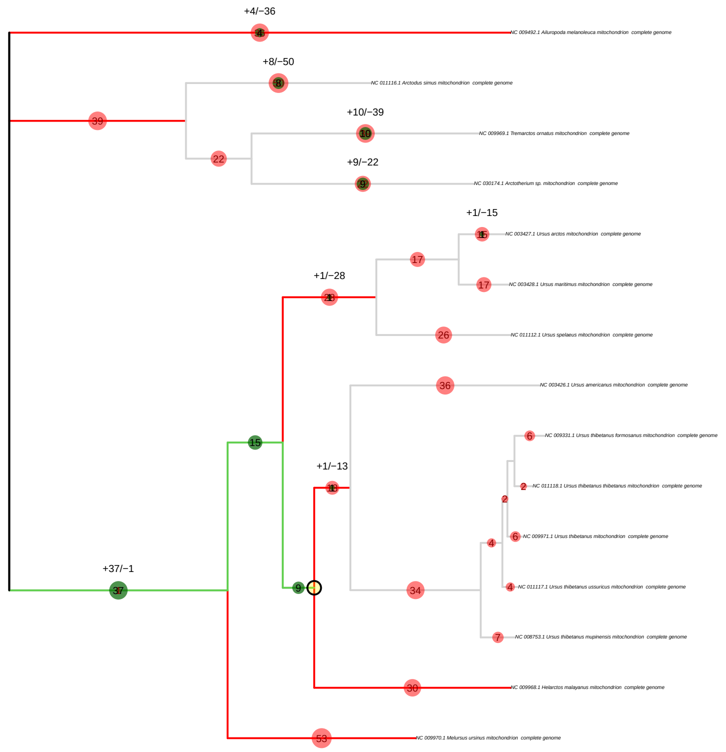


Figure 4: `pathPhynder` best path method results for the simulated ancient single-source sample N4 at $\sim 0.2X$ coverage. `pathPhynder` shows the best path for the given sample, which ends at the correct ancestral node N4.

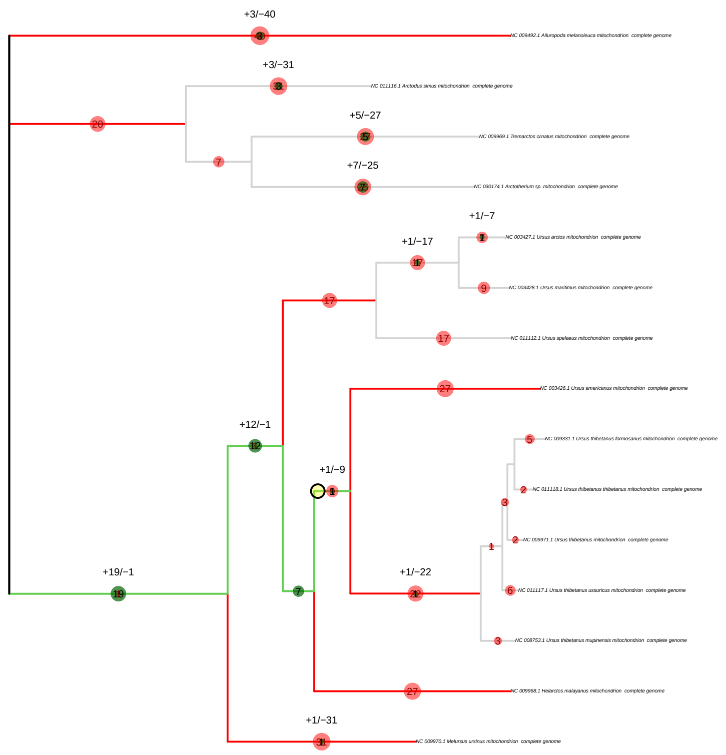


Figure 5: `pathPhynder` best path method results for the simulated ancient single-source sample N4 at $\sim 0.13X$ coverage. `pathPhynder` shows the best path for the given sample, which ends at one ancestral node after the target one N4 at N5.

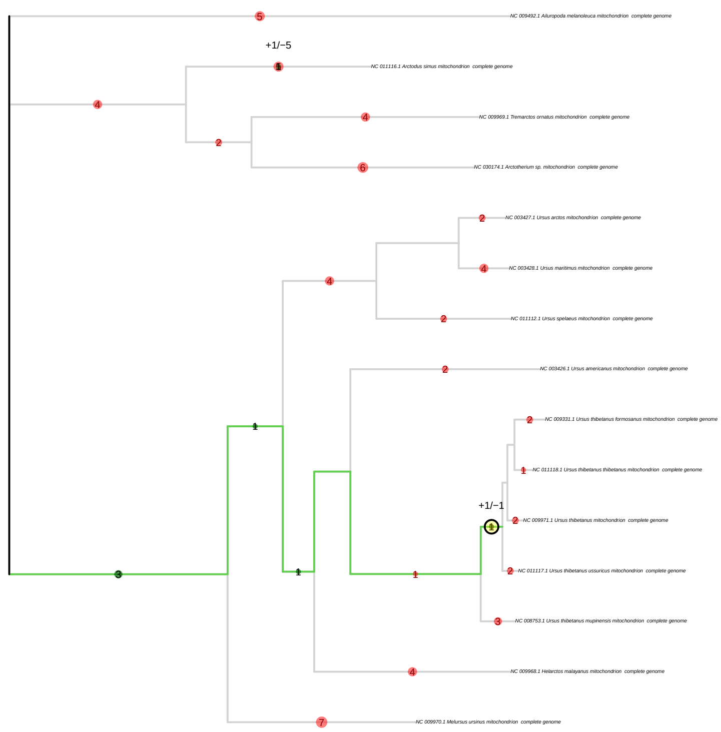


Figure 6: `pathPhynder` best path method results for the simulated ancient single-source sample N4 at $\sim 0.026X$ coverage. `pathPhynder` shows the best path for the given sample, which ends at the ancestral node N6.

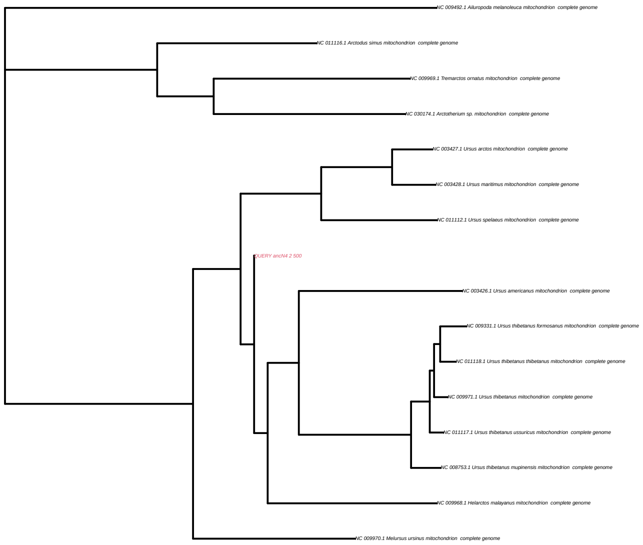


Figure 7: `pathPhynder` maximum likelihood method results for the simulated ancient single-source sample N4 at $\sim 1.3x$ coverage. `pathPhynder` shows the best path for the given sample, which ends at the correct ancestral node N4.

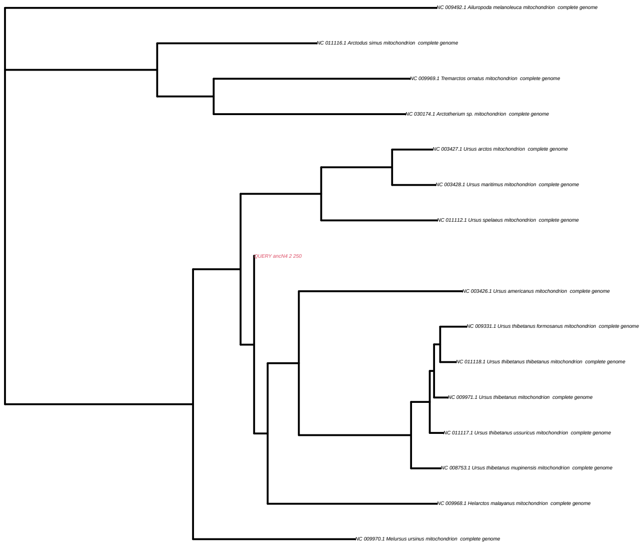


Figure 8: `pathPhynder` maximum likelihood method results for the simulated ancient single-source sample N4 at $\sim 0.66X$ coverage. `pathPhynder` shows the best path for the given sample, which ends at the correct ancestral node N4.

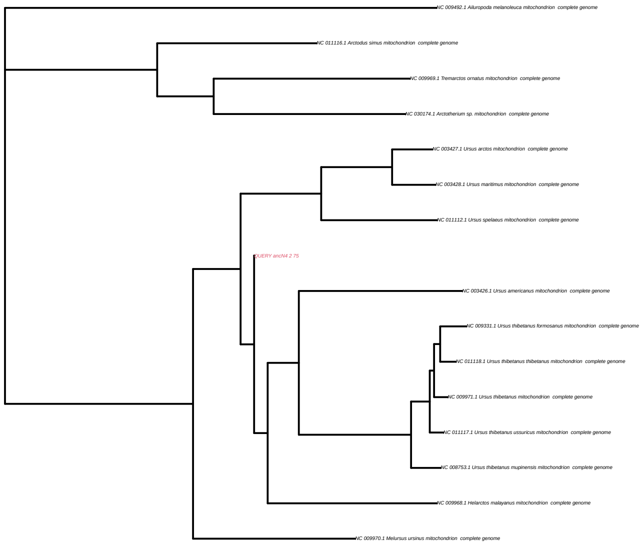


Figure 9: `pathPhynder` maximum likelihood method results for the simulated ancient single-source sample N4 at $\sim 0.2X$ coverage. `pathPhynder` shows the best path for the given sample, which ends at the correct ancestral node N4.

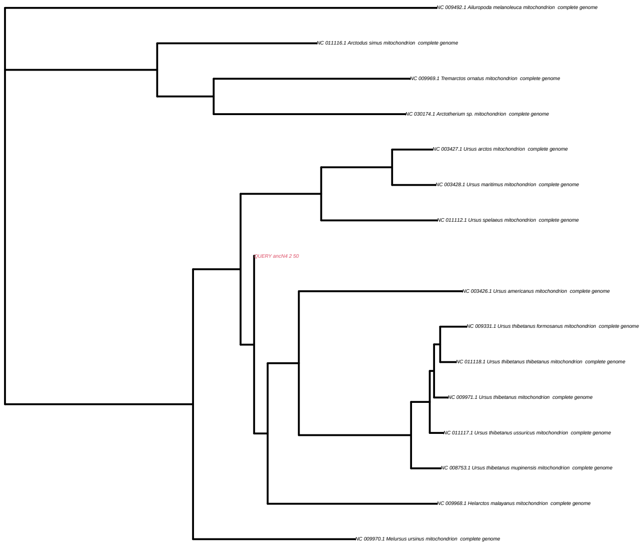


Figure 10: `pathPhynder` maximum likelihood method results for the simulated ancient single-source sample N4 at $\sim 0.13X$ coverage. `pathPhynder` shows the best path for the given sample, which ends at the correct ancestral node N4.

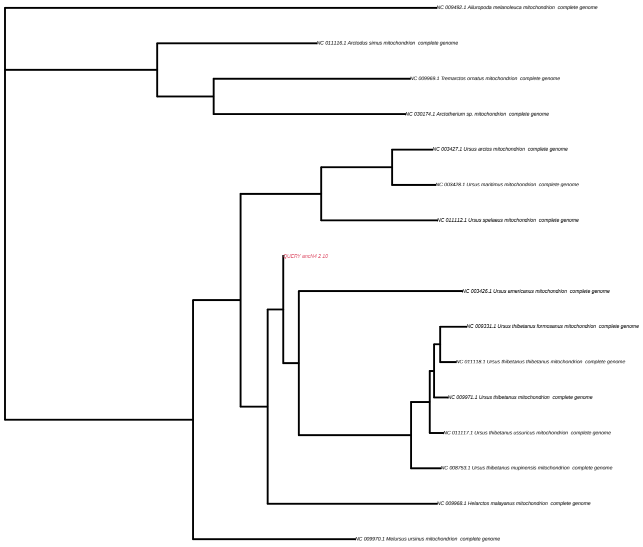


Figure 11: `pathPhynder` maximum likelihood method results for the simulated ancient single-source sample N4 at $\sim 0.026X$ coverage. `pathPhynder` shows the best path for the given sample, which ends at the ancestral node N6.

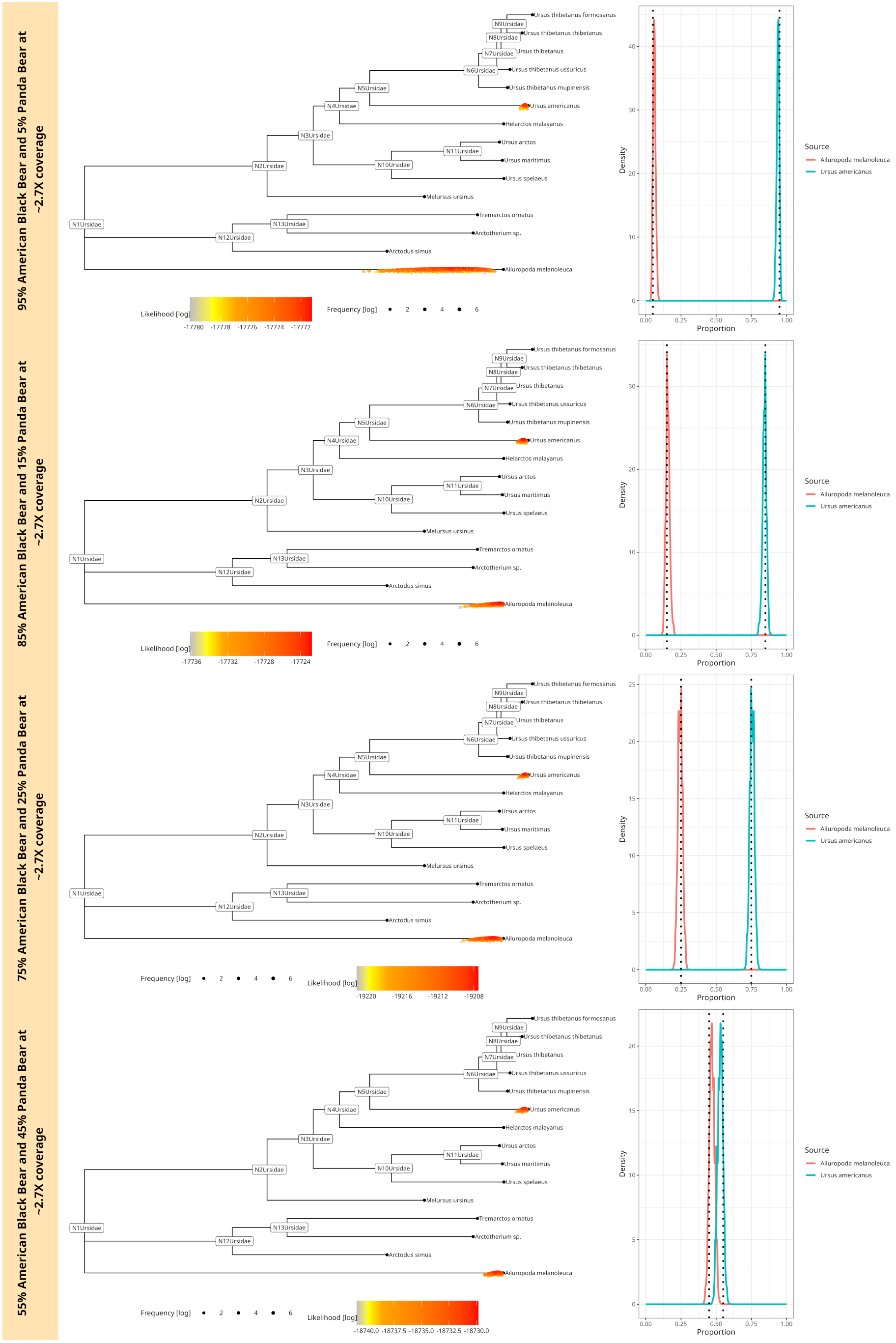


Figure 12: `soibean` results for simulated ancient fragments of two diverged species (83.4% similarity) from the family bears (Ursidae) at $\sim 2.7X$ coverage. The plot shows four different mixtures at $55\% - 45\%, 75\% - 25\%, 85\% - 15\%$ and $95\% - 5\%$ of the American Black bear and Giant Panda bear. The corresponding phylogenetic trees are displayed on the left: we plotted every accepted MCMC move coloured by likelihood value on the tree. The accepted moves are positioned above or below the tree, corresponding to a higher or lower likelihood value than the median, respectively. Each neighbouring plot shows the posterior proportion distribution, including the simulated proportion with a black dotted line.

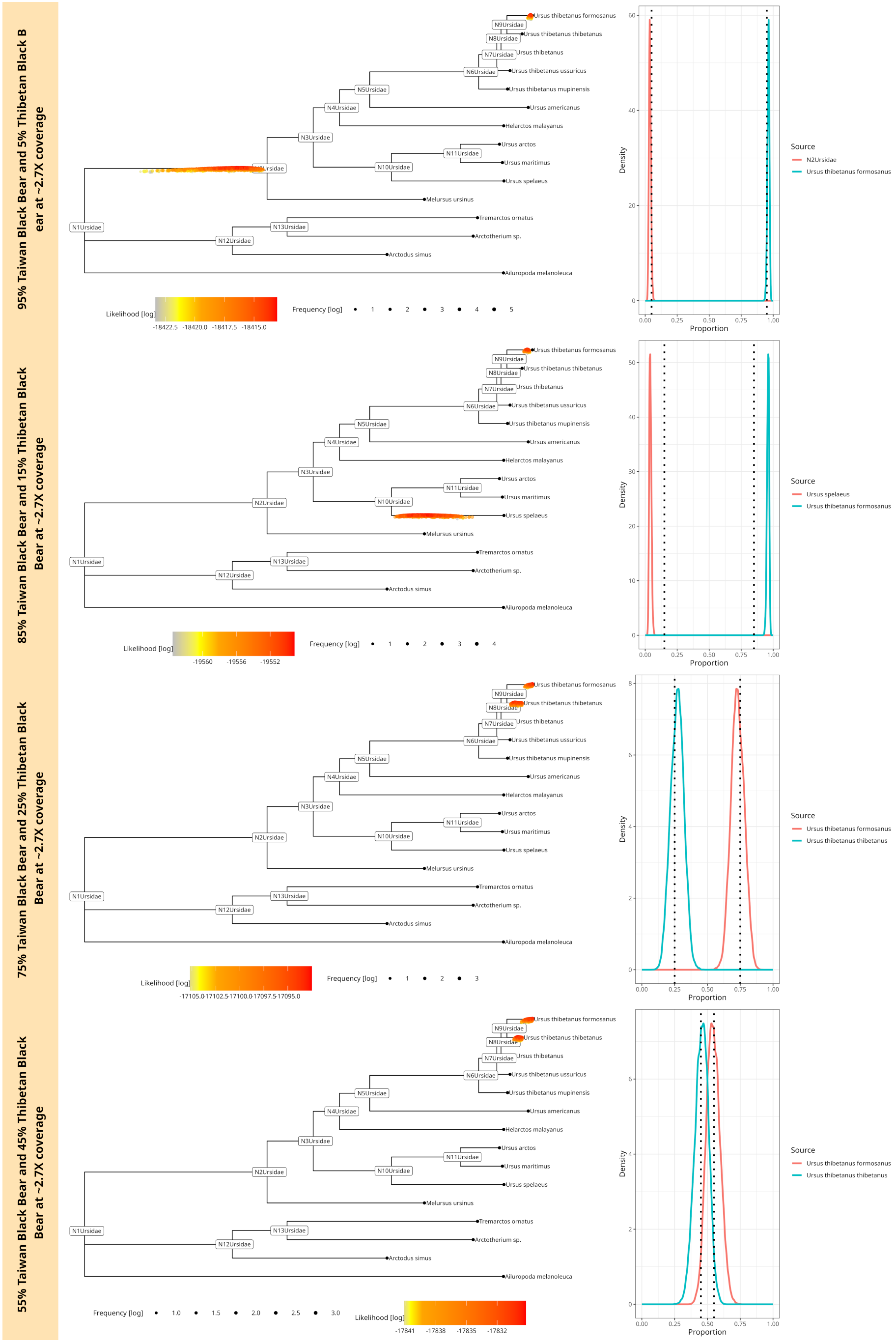


Figure 13: `soibean` results for simulated ancient fragments of two closely related species (98.8% similarity) from the family bears (Ursidae) at $\sim 2.7X$ coverage. The plot shows four different mixtures at $55\% - 45\%, 75\% - 25\%, 85\% - 15\%$ and $95\% - 5\%$ of the Taiwan and Tibetan Black bear. The corresponding phylogenetic trees are displayed on the left: we plotted every accepted MCMC move coloured by likelihood value on the tree. The accepted moves are positioned above or below the tree, corresponding to a higher or lower likelihood value than the median, respectively. Each neighbouring plot shows the posterior proportion distribution, including the simulated proportion with a black dotted line

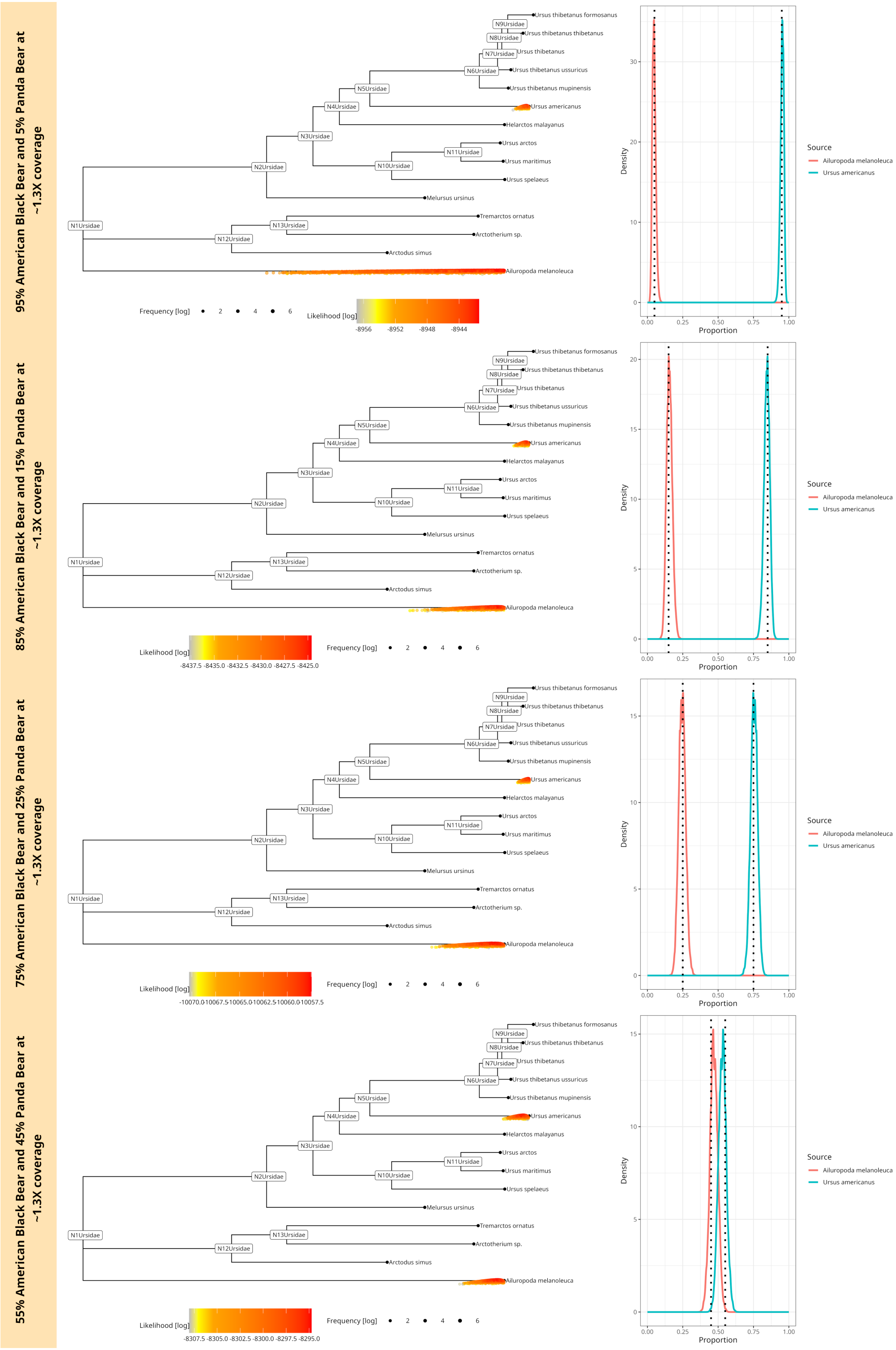


Figure 14: Mixtures of an American Black bear and a Giant Panda bear downsampled to $\sim 1.3X$ coverage (500 aDNA fragments). The plot shows four different mixtures at $55\% - 45\%, 75\% - 25\%, 85\% - 15\%$ and $95\% - 5\%$. The phylogenetic tree has a coloured point for each accepted MCMC move. The colour corresponds to the log-likelihood value. The posterior proportion distribution, including the simulated true proportion (black dotted line), is plotted on the right.

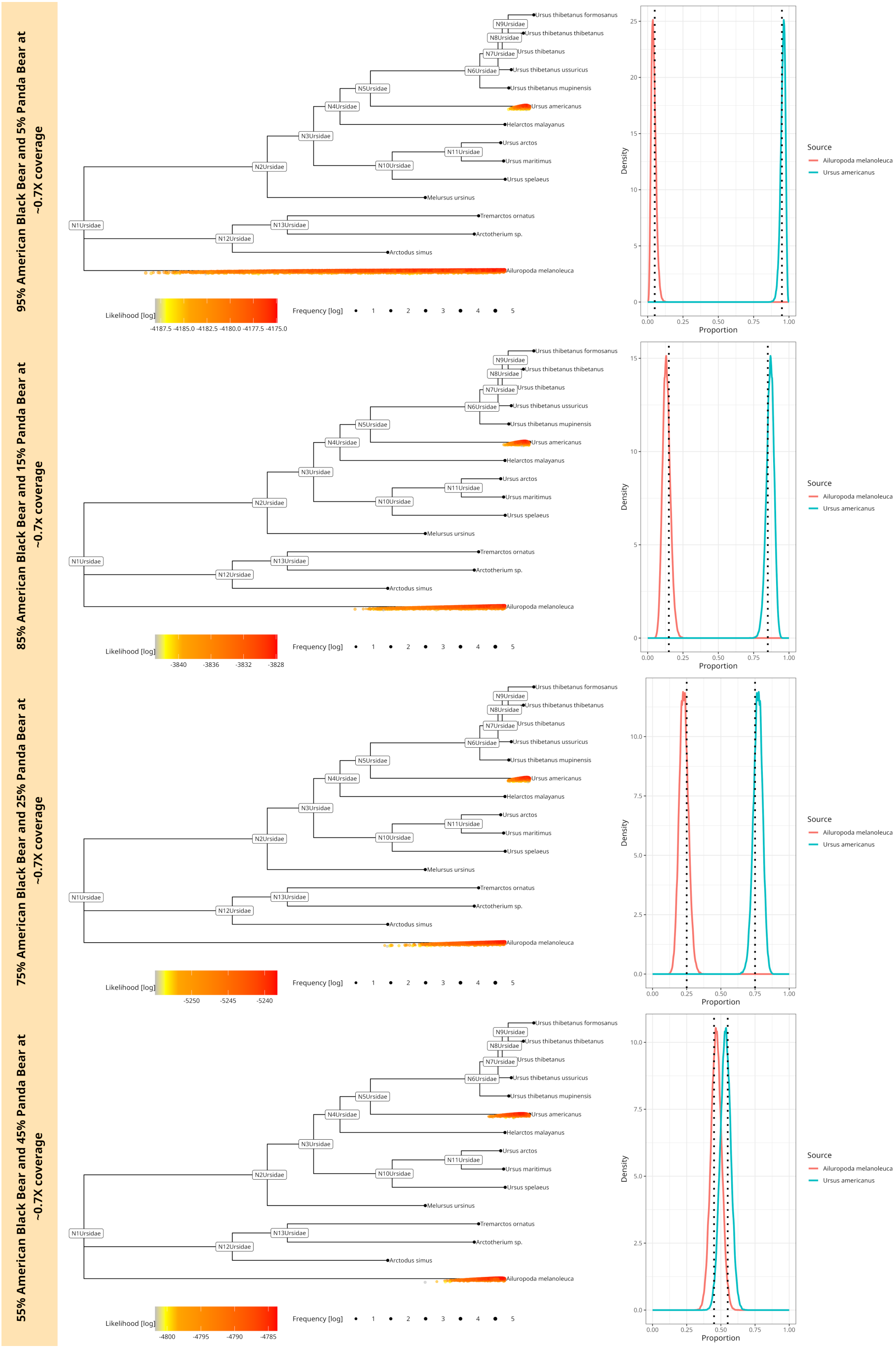


Figure 15: Mixtures of an American Black bear and a Giant Panda bear downsampled to $\sim 0.7x$ coverage (250 aDNA fragments). The plot shows four different mixtures at $55\% - 45\%, 75\% - 25\%, 85\% - 15\%$ and $95\% - 5\%$. The phylogenetic tree has a coloured point for each accepted MCMC move. The colour corresponds to the log-likelihood value. The posterior proportion distribution, including the simulated true proportion (black dotted line), is plotted on the right.

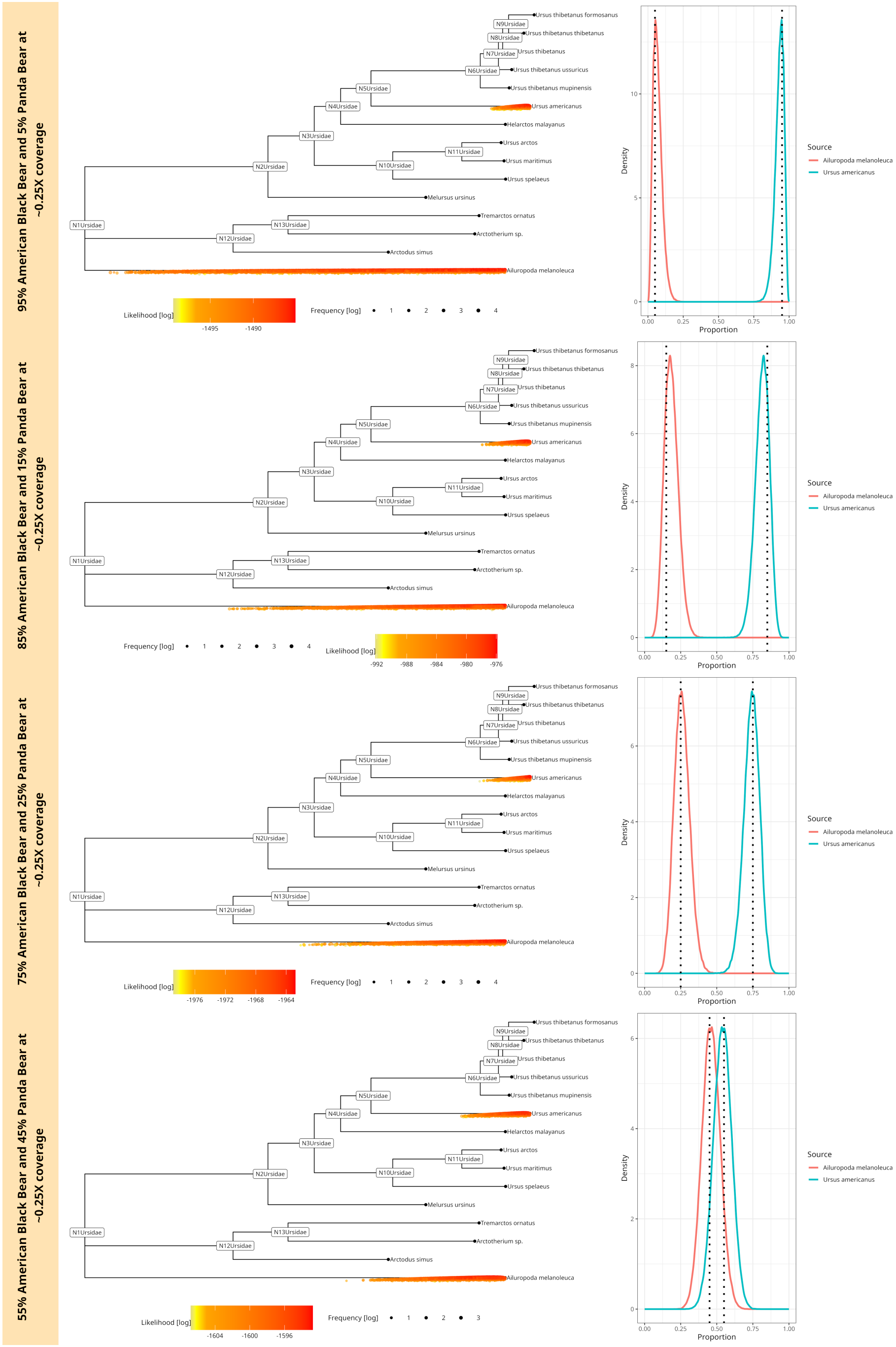


Figure 16: Mixtures of an American Black bear and a Giant Panda bear downsampled to $\sim 0.25x$ coverage (100 aDNA fragments). The plot shows four different mixtures at $55\% - 45\%, 75\% - 25\%, 85\% - 15\%$ and $95\% - 5\%$. The phylogenetic tree has a coloured point for each accepted MCMC move. The colour corresponds to the log-likelihood value. The posterior proportion distribution, including the simulated true proportion (black dotted line), is plotted on the right.

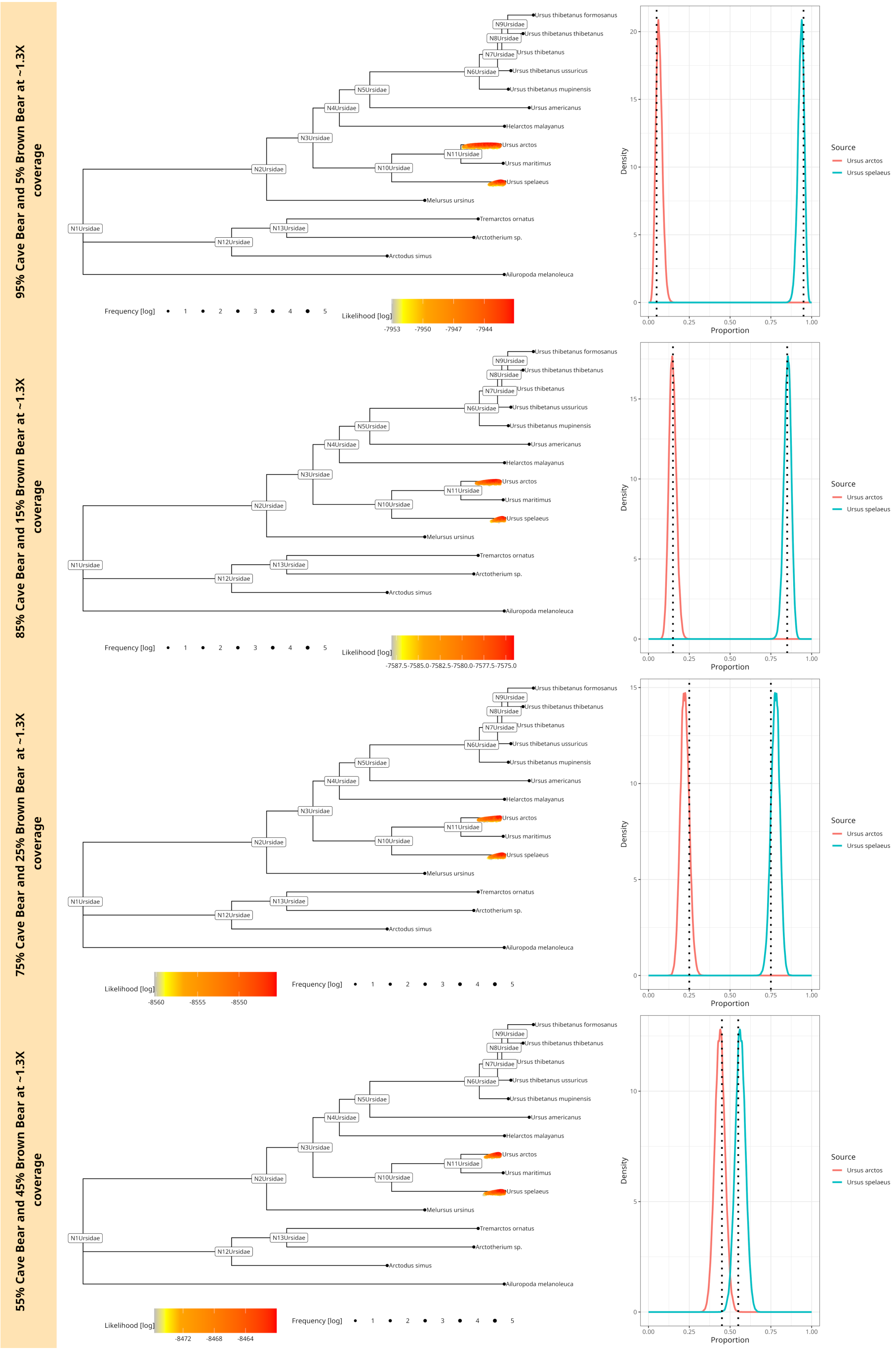


Figure 17: Cave bear and Brown bear mixture downsampled to $\sim 1.3X$ coverage (500 aDNA fragments). The plot shows four different mixtures at $55\% - 45\%, 75\% - 25\%, 85\% - 15\%$ and $95\% - 5\%$. The phylogenetic tree has a coloured point for each accepted MCMC move. The colour corresponds to the log-likelihood value. The posterior proportion distribution, including the simulated true proportion (black dotted line), is plotted on the right.

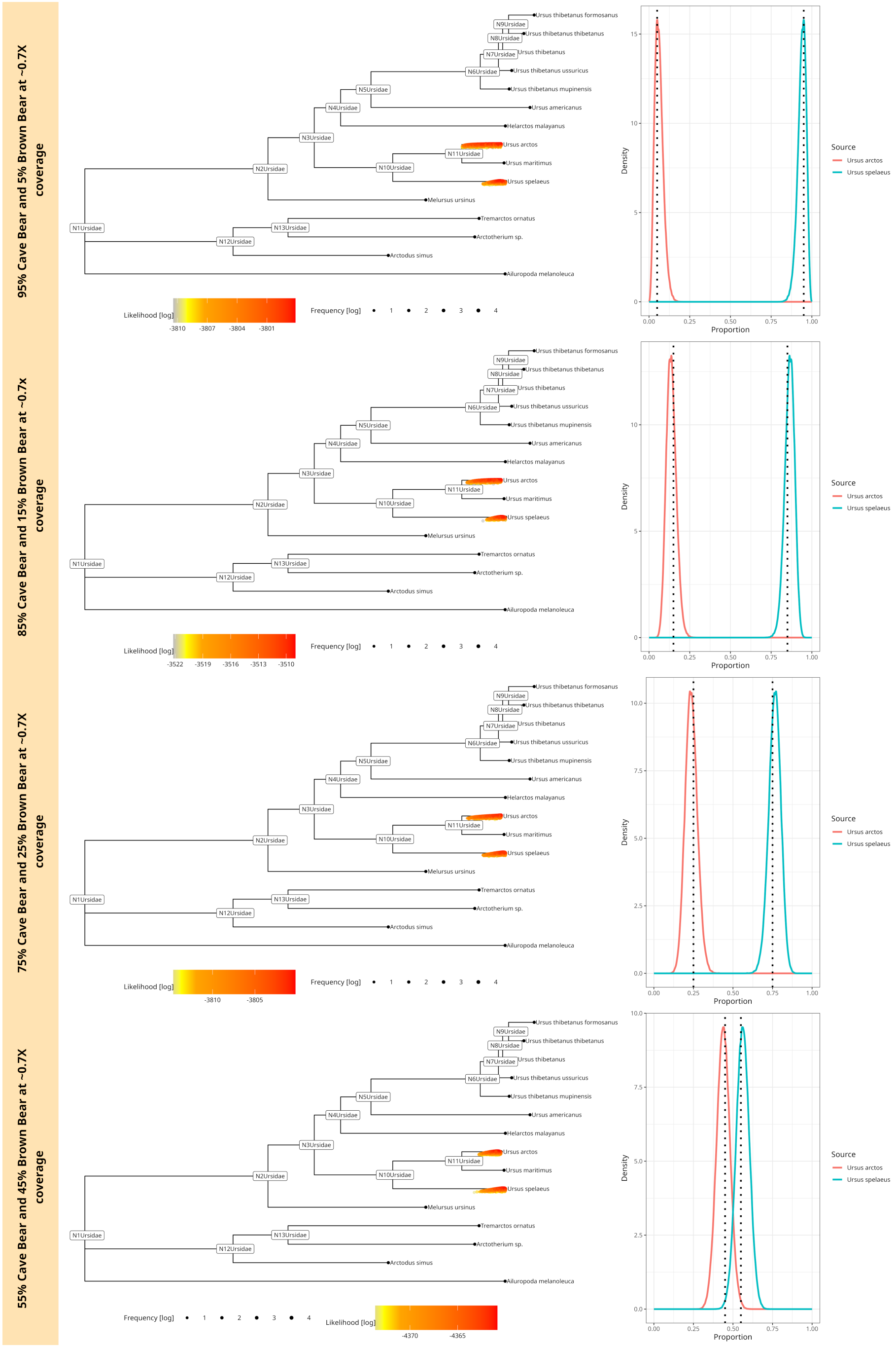


Figure 18: Cave bear and Brown bear mixture downsampled to $\sim 0.7X$ coverage (250 aDNA fragments). The plot shows four different mixtures at $55\% - 45\%, 75\% - 25\%, 85\% - 15\%$ and $95\% - 5\%$. The phylogenetic tree has a coloured point for each accepted MCMC move. The colour corresponds to the log-likelihood value. The posterior proportion distribution, including the simulated true proportion (black dotted line), is plotted on the right.

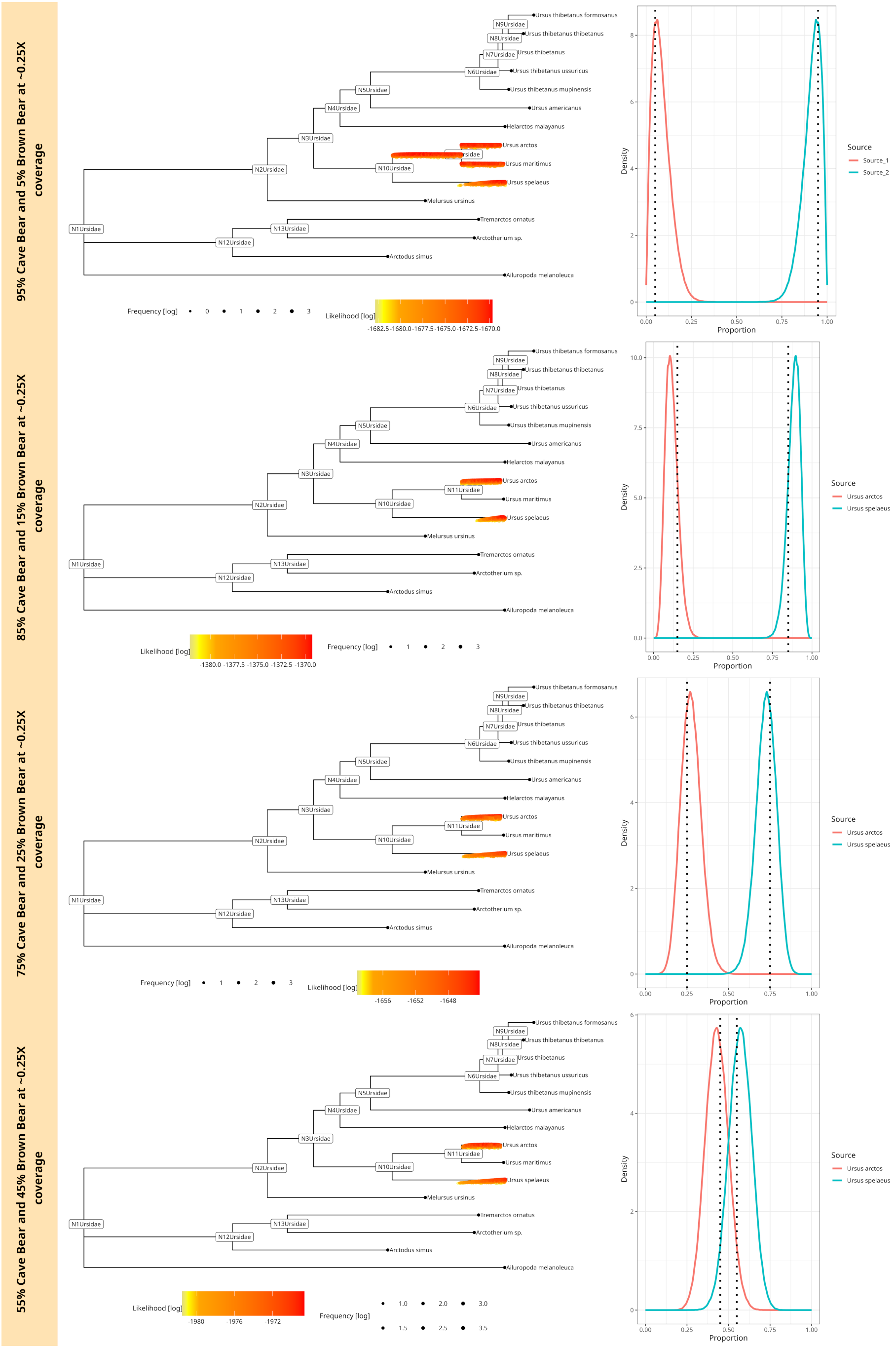


Figure 19: Mixtures of a Cave bear and a Brown bear downsampled to $\sim 0.25X$ coverage (100 aDNA fragments). The plot shows four different mixtures at $55\% - 45\%, 75\% - 25\%, 85\% - 15\%$ and $95\% - 5\%$. The phylogenetic tree has a coloured point for each accepted MCMC move. The colour corresponds to the log-likelihood value. The posterior proportion distribution, including the simulated true proportion (black dotted line), is plotted on the right.

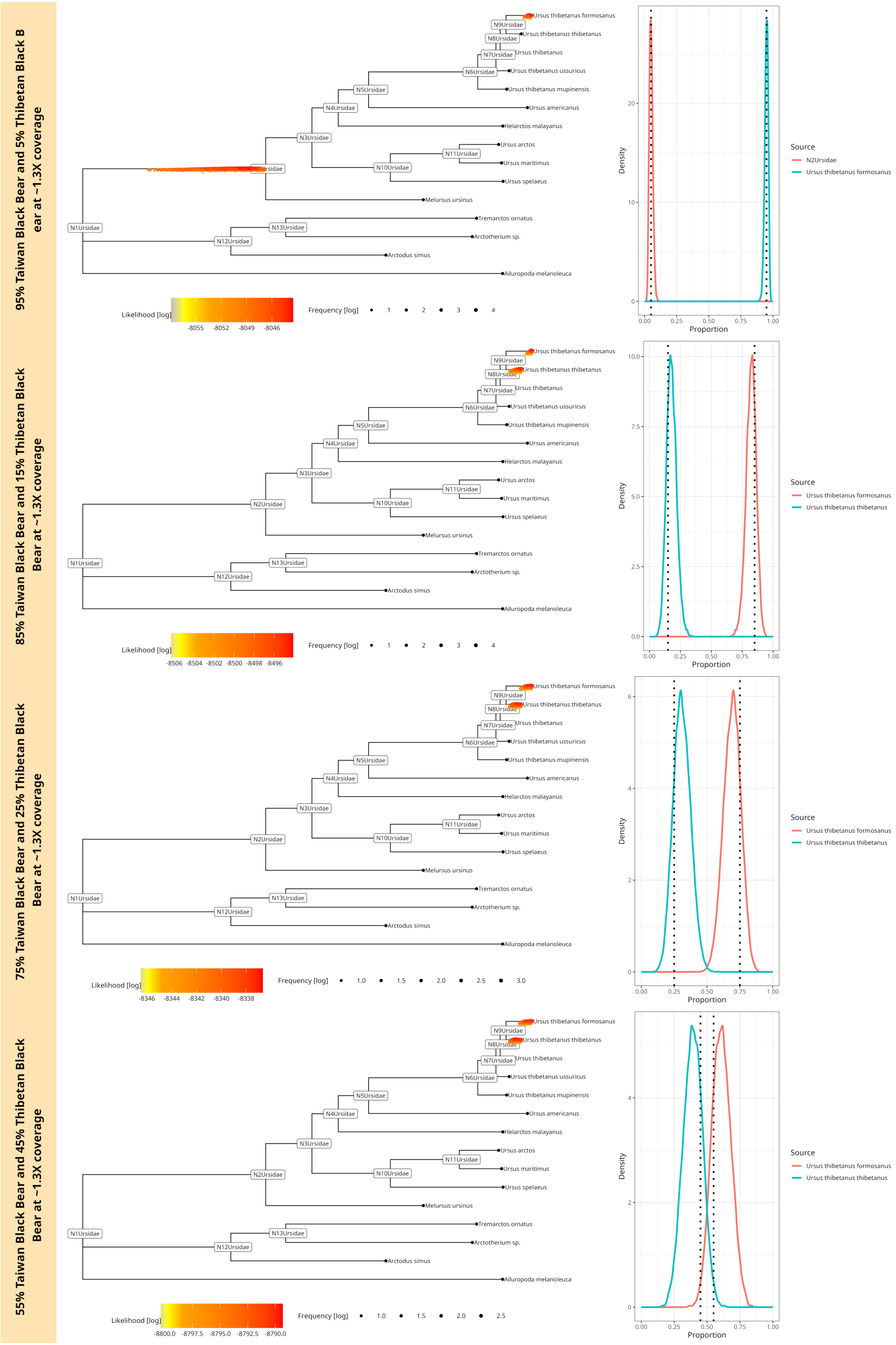


Figure 20: Mixtures of a Taiwan and Tibetan Black bear downsampled to $\sim 1.3X$ coverage (500 aDNA fragments). The plot shows four different mixtures at $55\% - 45\%, 75\% - 25\%, 85\% - 15\%$ and $95\% - 5\%$. The phylogenetic tree has a coloured point for each accepted MCMC move. The colour corresponds to the log-likelihood value. The posterior proportion distribution, including the simulated true proportion (black dotted line), is plotted on the right.

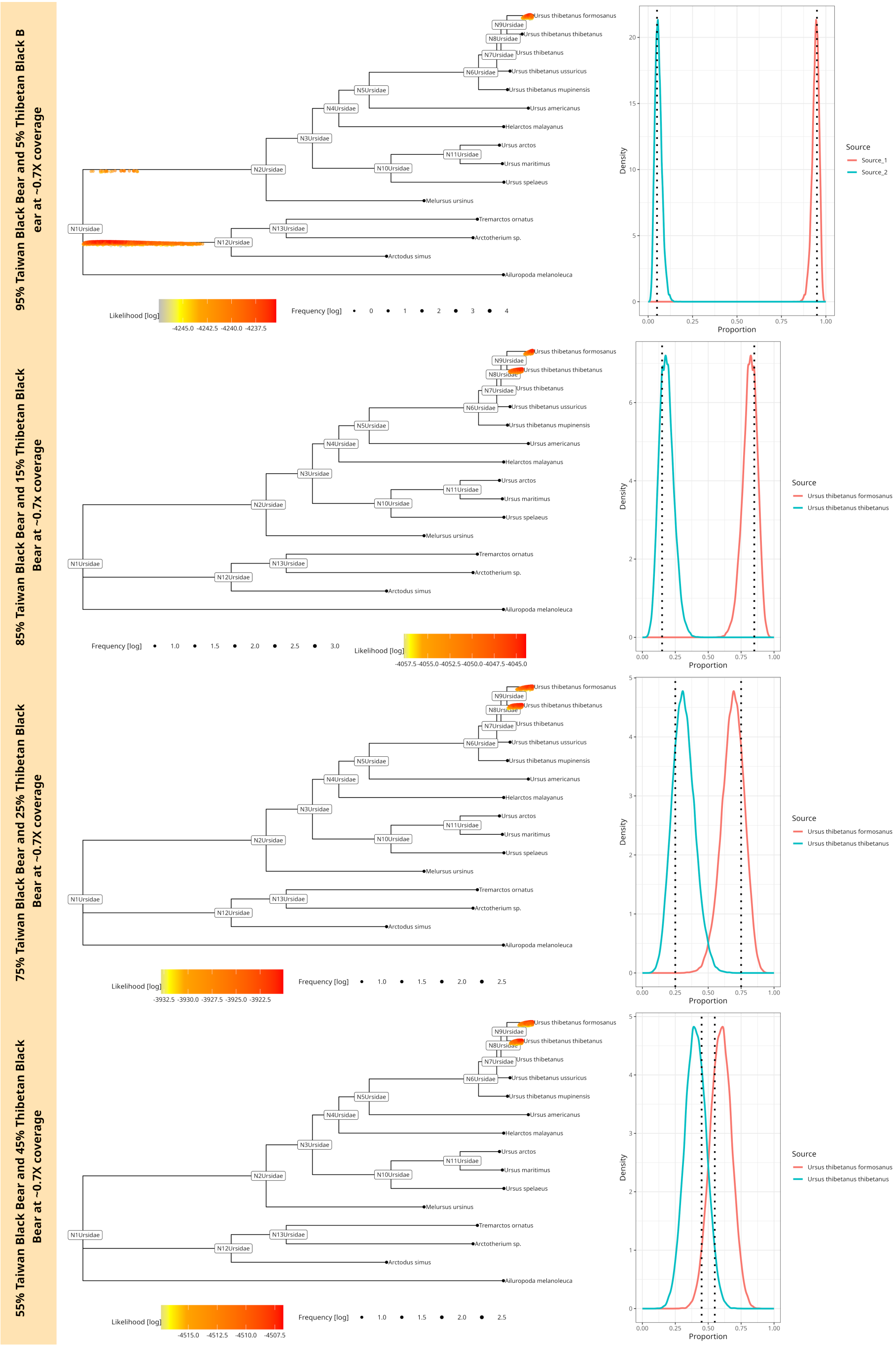


Figure 21: Mixtures of a Taiwan and Tibetan Black bear downsampled to $\sim 0.7X$ coverage (250 aDNA fragments). The plot shows four different mixtures at $55\% - 45\%, 75\% - 25\%, 85\% - 15\%$ and $95\% - 5\%$. The phylogenetic tree has a coloured point for each accepted MCMC move. The colour corresponds to the log-likelihood value. The posterior proportion distribution, including the simulated true proportion (black dotted line), is plotted on the right.

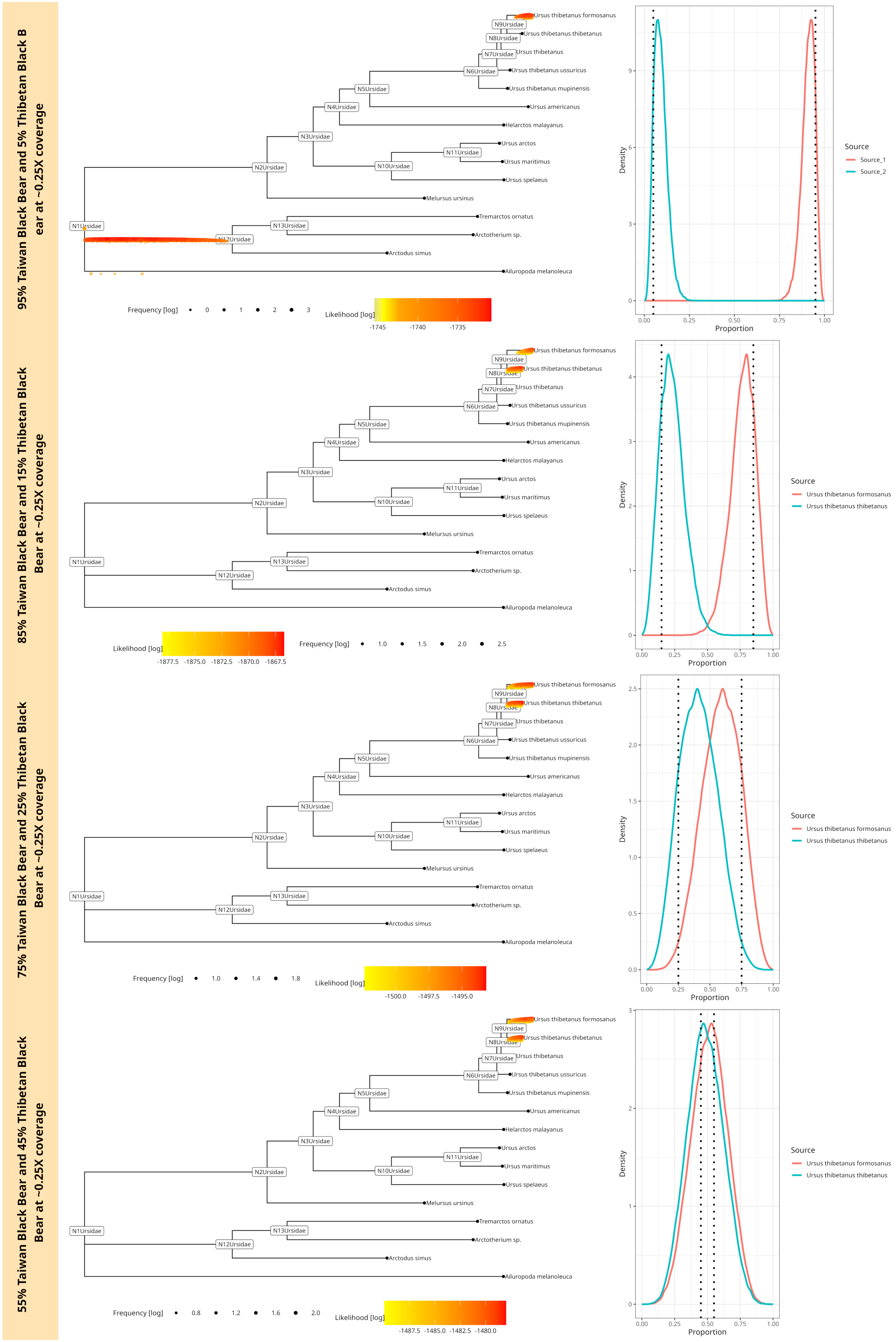

Figure 22: Mixtures of a Taiwan and Tibetan Black bear downsampled to $\sim 0.25X$ coverage (100 aDNA fragments). The plot shows four different mixtures at $55\%-45\%, 75\%-25\%, 85\%-15\%$ and $95\%-5\%$. The phylogenetic tree has a coloured point for each accepted MCMC move. The colour corresponds to the log-likelihood value. The posterior proportion distribution, including the simulated true proportion (black dotted line), is plotted on the right.

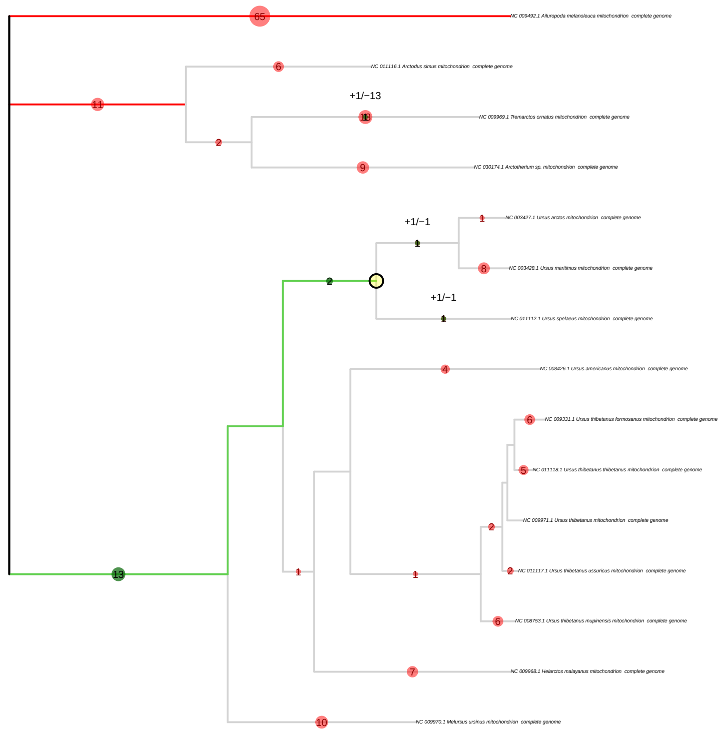


Figure 23: `pathPhynder` results for the simulated ancient mixture of a cave bear (*Ursus spelaeus*) 55% and the brown bear (*Ursus arctos*) 45% at $\sim 2.7X$ coverage. `pathPhynder` shows the best path for the given sample, which ends at the lowest common ancestor for the given mixture.

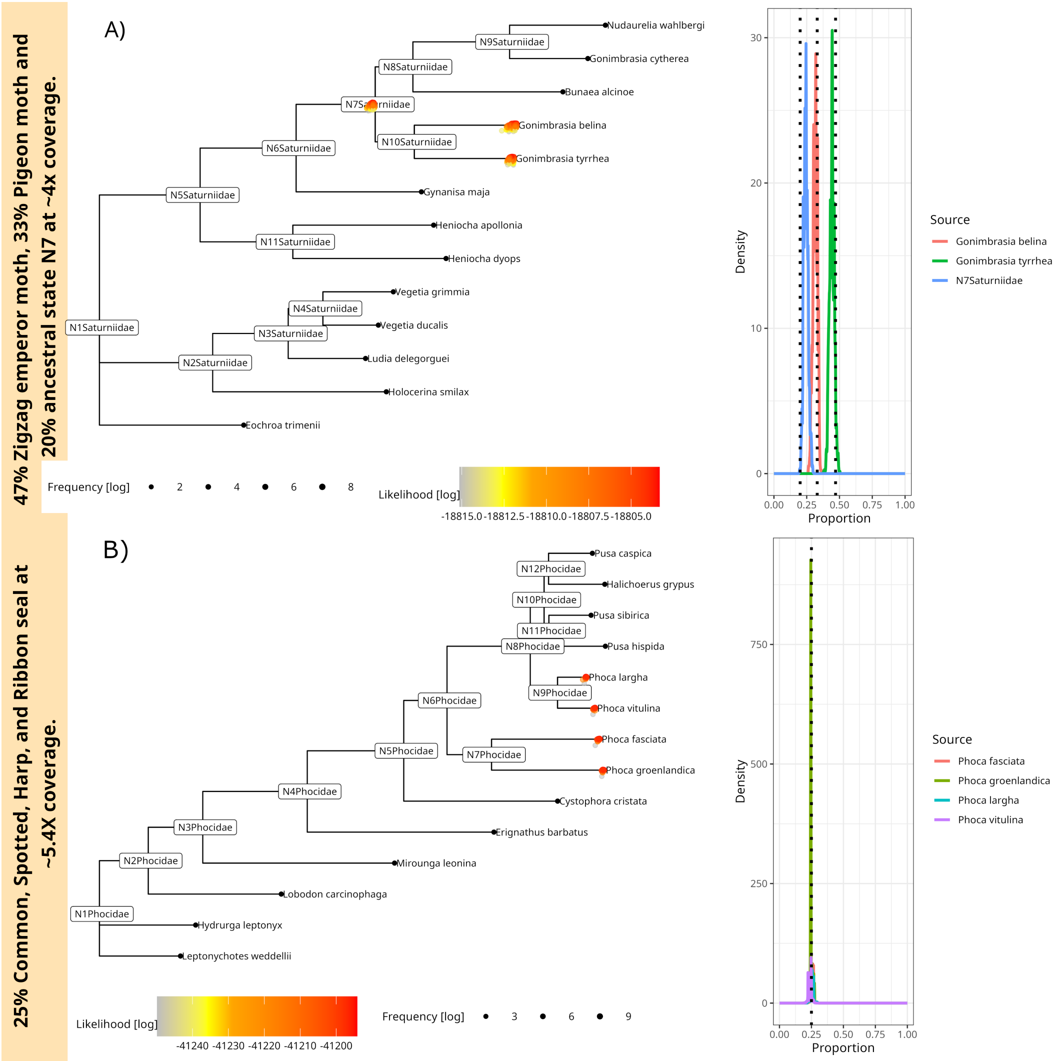


Figure 24: A) $44\% - 30\% - 23\%$ mixture of a three-source simulated ancient sample from a family of winged insects (Saturniidae). The mixture contains two emperor moths (*Gonimbrasia tyrrhea* and *Gonimbrasia belina* as well as the ancestral state N7. We sampled to$\sim 4X$ coverage. The tree shows every accepted MCMC move, coloured and placed by log-likelihood value, where a position above the tree branch represents a better likelihood than the median likelihood and a position underneath the tree branch a worse likelihood (right side). The right side of the plot shows `soibean`'s proportion estimation with the simulated true proportion represented with a black dotted line. B) $25\% - 25\% - 25\% - 25\%$ mixture of a four-source simulated ancient sample from the family of seals (Phocidae). The mixture contains the earless seal species, namely *Phoca largha*, *Phoca vitulina*, *Phoca groenlandica* and *Pusa hispida* at $\sim 5.4X$ coverage (2000 aDNA fragments). The tree shows every accepted MCMC move, coloured and placed by log-likelihood value plot, showing `soibean`'s proportion estimation with the simulated true proportion with a black dotted line.

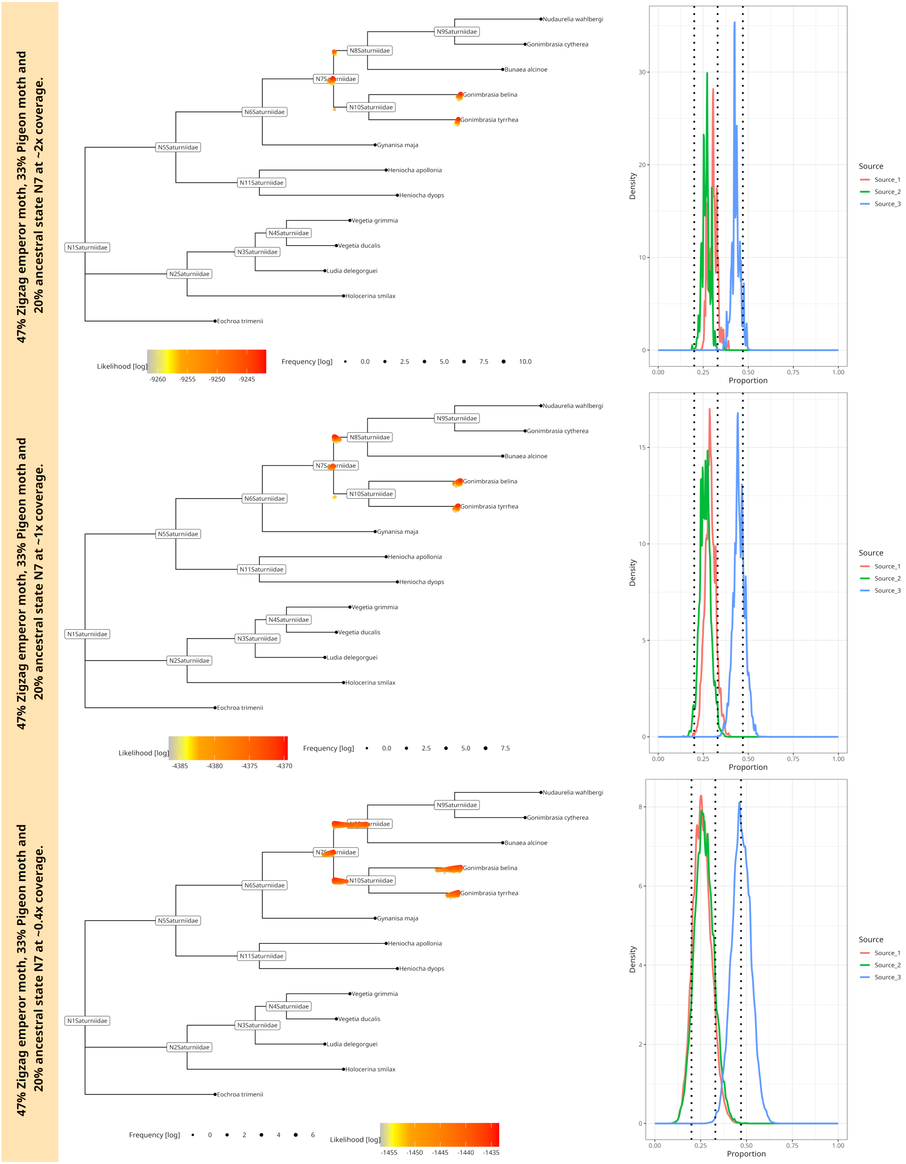


Figure 25: $44\% - 30\% - 23\%$ mixture of a three-source simulated ancient sample from a family of winged insects (Saturniidae). The mixture contains two emperor moths (*Gonimbrasia tyrrhea* and *Gonimbrasia belina* as well as the ancestral state N7 and is downsampled to $\sim 2X$ coverage (750 aDNA fragments), $\sim 1X$ coverage (375 aDNA fragments) and $\sim 0.4x$ coverage (150 aDNA fragments). The trees show every accepted MCMC move, coloured and placed by log-likelihood value (left side) and `soibean`'s proportion estimations on the left side, where the black dotted line shows the simulated proportion.

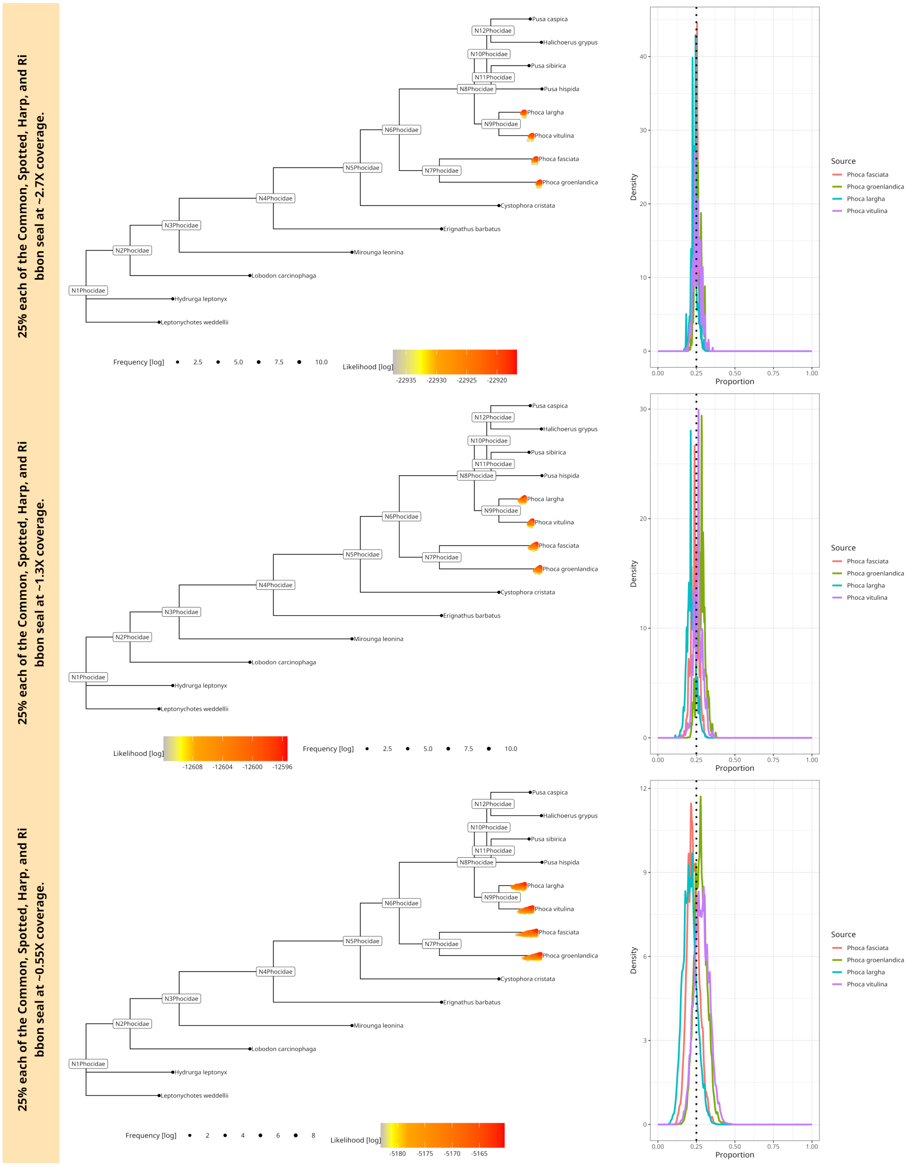

Figure 26: 25% – 25% – 25% – 25% mixture of a four-source simulated ancient sample from the family of seals (Phocidae). The mixture contains the earless seal species, namely *Phoca largha*, *Phoca vitulina*, *Phoca groenlandica* and *Pusa hispida*; it is downsampled to $\sim 2.7X$ coverage (1000 aDNA fragments), $\sim 1.3X$ coverage (500 aDNA fragments) and $\sim 0.55X$ coverage (200 aDNA fragments). The trees show every accepted MCMC move, coloured and placed by log-likelihood value (left side) and `soibean`'s proportion estimations on the left side, where the black dotted line shows the simulated proportion.

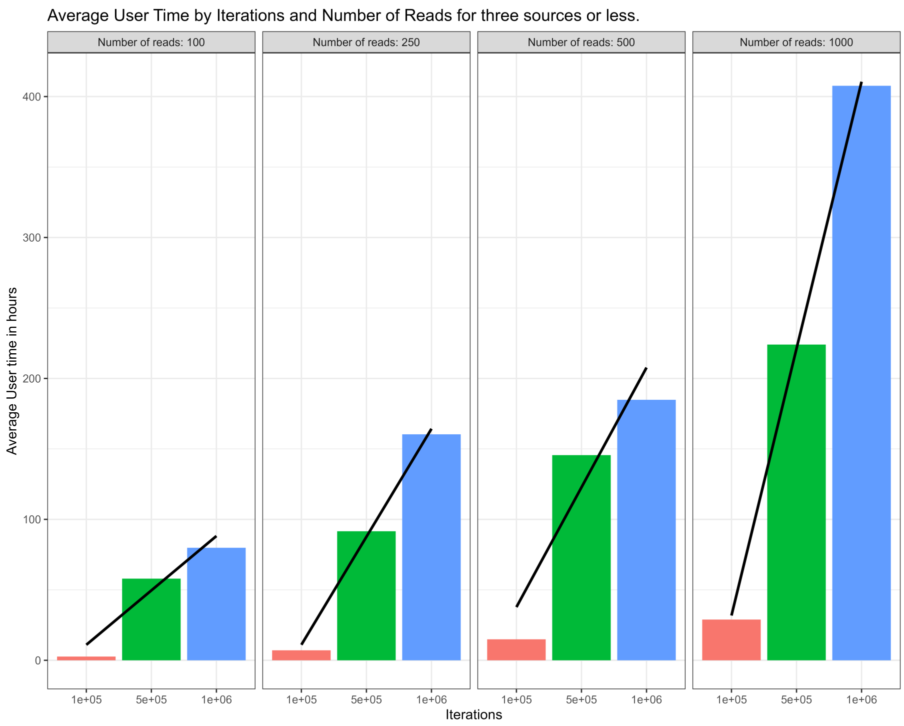


Figure 27: `soibean` user-time (hours) comparison for three different numbers of reads and three different numbers of iterations.

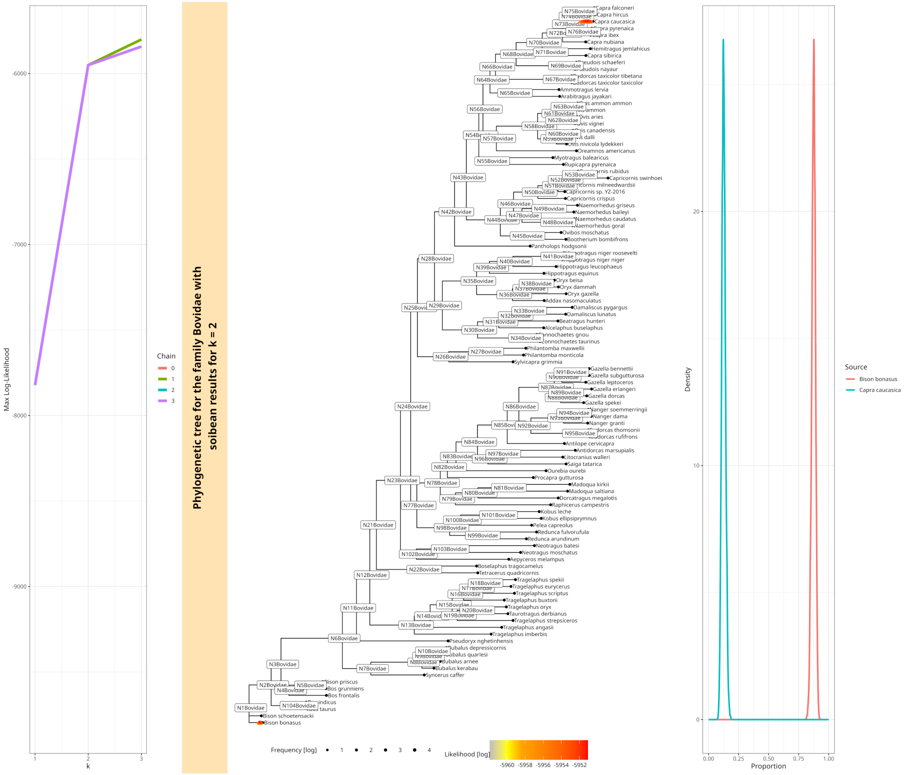


Figure 28: The results from `soibean` for the 25kya sediment sample from a Georgian cave. The k-curve highlights the presence of two contributing sources for the filtered Bovidae fragments. Analyzing the results, we can find the previously detected European Bison (*Bison bonasus*). Additionally, we can identify a low-coverage sample from the West Caucasian tur (*Capra caucasica*, with a proportion of about 10%, which was previously not reported.

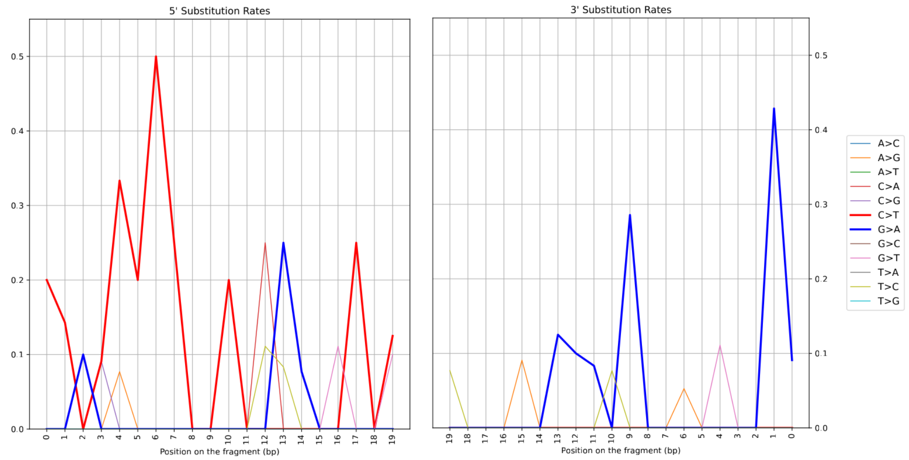


Figure 29: Deamination patterns for the alignments to the mitochondrial genome of *Capra caucasica*.

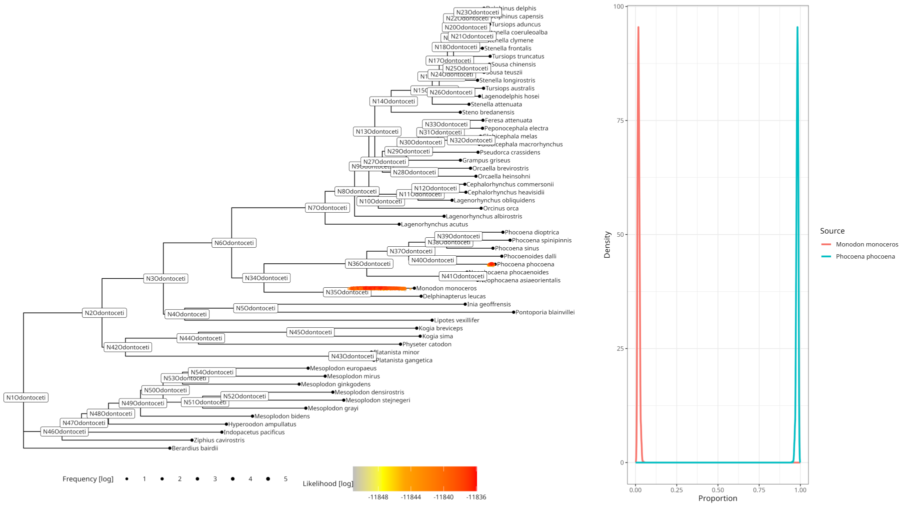


Figure 30: `soibean` results for the 9500-year-old empirical metagenomic sample from pitch pieces found in Huseby Klev on the northwestern coast of Sweden for $k = 3$. The harbour porpoise (*Phocoena phocoena*) is still the most likely source. `soibean` distributes one source at about 5% to an ancestral state of two Asiatic river dolphins. `soibean`'s branch and source proportion diagnostics showed an effective sample size below 200, indicating uncertainty for the second source.

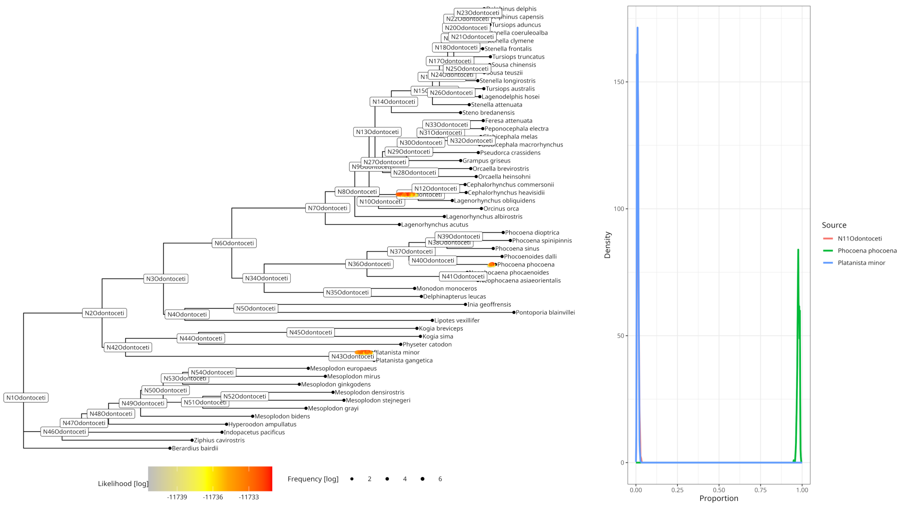


Figure 31: `soibean` results for the 9500-year-old empirical metagenomic sample from pitch pieces found in Huseby Klev on the northwestern coast of Sweden for $k = 3$. The harbour porpoise (*Phocoena phocoena*) is still the most likely source. `soibean` distributes two sources at about 2.5% to an ancestral state of two Asiatic river dolphins and a third dolphin species. Both additional sources have an effective sample size below 200 in `soibean`'s branch and source proportion diagnostics, making them unlikely contributing sources.

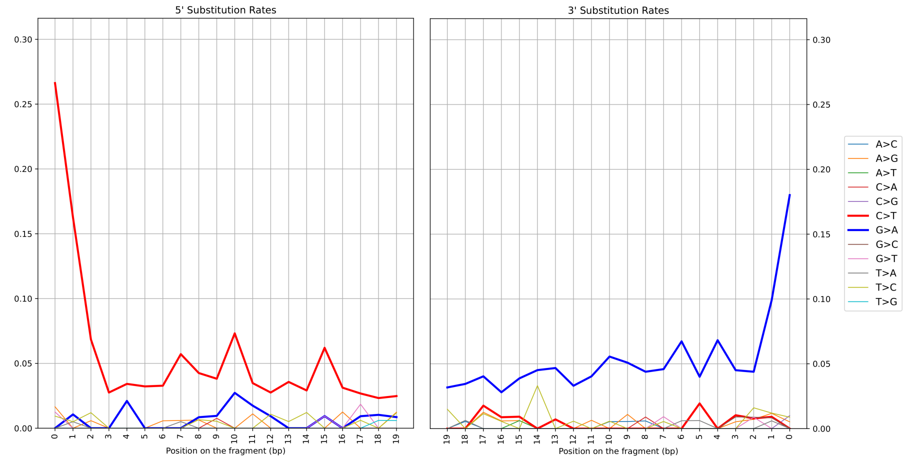


Figure 32: Complete deamination patterns for the alignments to the mitochondrial genome of *Phocoen*
*phocoena*.
